# Hierarchical Temporal Transformer for Cancer Grade Prediction and Cross-Cancer Transfer Learning from Pathology Reports

**DOI:** 10.64898/2026.11.05.764680

**Authors:** Nura Brimo, Rajneesh Anand, Anshuman Sabath, Hassan Harb, Dilek Çökeliler Serdaroğlu

## Abstract

**Background:** Language models for cancer reports usually read one document at a time and are tested on tumour types seen in training. Continuous-time sequence models address the first gap, but whether they use visit timing, not just order, is seldom tested.

**Methods:** We developed the Hierarchical Temporal Transformer (HTT), which encodes reports with adapted BiomedBERT and reads the sequence with a cancer-type-conditioned temporal transformer. We evaluated it against best-possible AUROC ceilings on two synthetic benchmarks, one with the label stated in the text and one recoverable only from tumour sizes and visit intervals (ten seeds). Diagnostics led to an interval-aware time code (HTT-IA). On 4,786 TCGA pathology reports in 14 types, three types were withheld, and a text-only HTT was compared with a survival model.

**Results:** On the phrase benchmark HTT reached AUROC 0.931 ± 0.012 against 0.913 ± 0.007 for a last-report baseline (+0.018, 95% CI −0.009 to +0.050); dates cannot raise the ceiling there. On the growth benchmark the initial time code scored 0.790, equal to a no-time model (0.787), whereas HTT-IA reached 0.843 (+0.053, 0.026 to 0.082) against ceilings of 0.894 without and 0.979 with dates. On unseen tumour types HTT reached 0.925 ± 0.007, equal to in-distribution performance and to a single-report baseline. Text-only macro C-index was 0.592 (CALM 0.626–0.652).

**Conclusion:** Cross-type transfer is carried by the pre-trained encoder. Visit timing is used only when the data make it identifiable and the time code is on a clinical scale. Ceilings, order-only ablations and re-dating probes are inexpensive checks for such models.

**Statement of significance:** 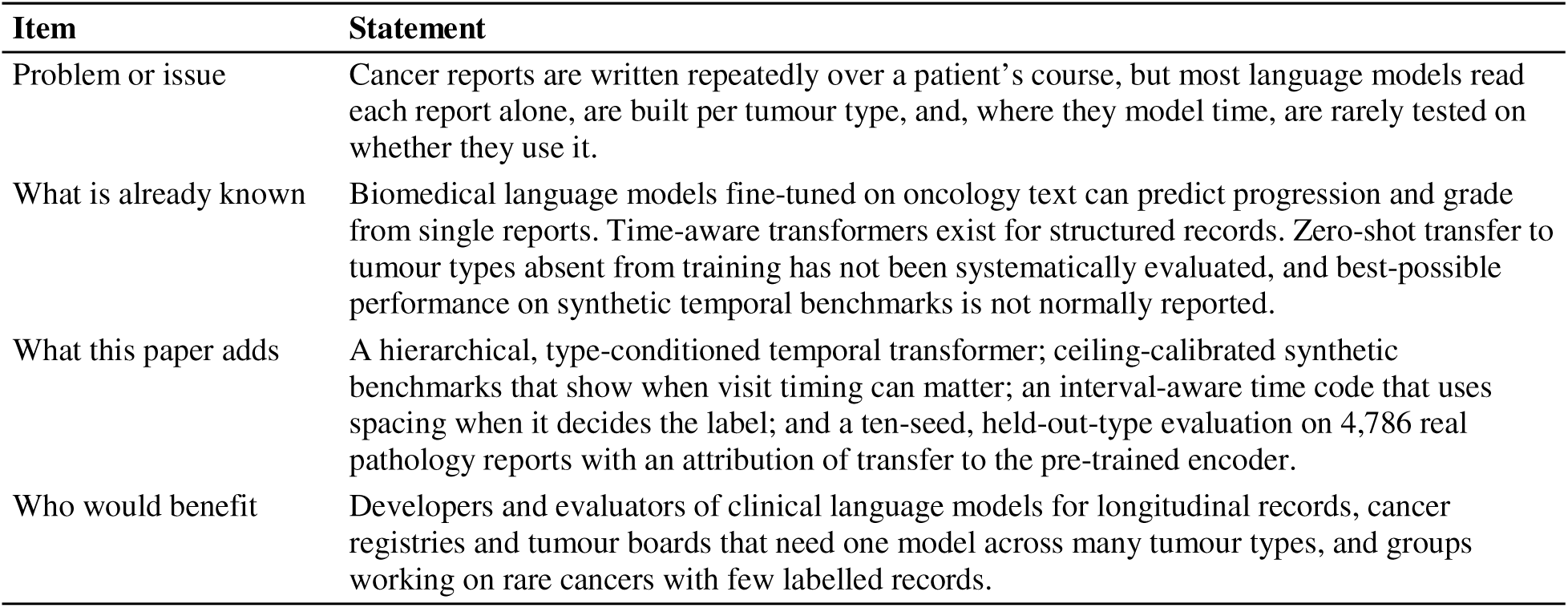

## 1. Introduction

Cancer care produces text at every stage. Radiologists write imaging impressions, pathologists describe resected tissue, and oncologists synthesise both in clinical notes [1]. Language models that read these documents have proved useful for predicting progression from radiology impressions [2,3], extracting phenotypes and generating diagnoses [4], and classifying disease response [5]. Like clinical language models more generally [6], they share two limitations that this study addresses.

The first is that each report is read in isolation. Patients are seen repeatedly over months or years, and the shape of that history carries information. A patient whose scans move from stable disease to enlargement to new nodal involvement is in a different situation from one whose latest scan alone mentions new spread, even when the final reports read identically; a model that sees only the last report cannot tell them apart. Timing adds a further layer. A tumour that has grown by thirty percent between two scans conveys one thing if the scans were three weeks apart and another if they were three months apart, and response criteria such as RECIST define change relative to a baseline without stating the rate that a clinician actually reasons about [7]. Models of longitudinal health records have long taken time into account, from time-aware recurrent networks [8] to transformers pretrained on visit sequences [9,10] (reviewed in [11]). For transformers the natural device is a continuous-time positional encoding, in which the integer position of the original architecture [12] is replaced by a function of a real-valued timestamp [13].

The second limitation is the absence of cross-cancer-type transfer. A comprehensive cancer centre treats dozens of tumour types, and building and maintaining a separate model for each scales poorly. For rare cancers the problem is feasibility rather than cost, since an institution may never accumulate enough labelled records to train at all [14]. Prior work either trains and tests on a single type or, where several types are used, evaluates only on types seen during training. Whether a model trained on breast, lung and kidney cancer can read a thyroid or sarcoma report it has never encountered is, to our knowledge, not systematically known.

A third issue concerns how temporal models are evaluated, and it shapes the design of this study. Whether a model uses timing, as opposed to merely accepting timestamps as input, depends on whether the data make timing informative. On a benchmark in which the label is written into the text and the dates are unrelated to it, a model that ignores the dates is behaving optimally, and no result on that benchmark can show otherwise. Synthetic benchmarks are attractive because their data-generating process is known, so the best-possible performance of a reader with and without the timing information can be computed. Combined with intervention on the timing input, and with enough training seeds and test patients to see effects of the claimed size [15], this turns ‘does the model use time?’ into a question with an answer.

We developed the Hierarchical Temporal Transformer (HTT) to address the first two limitations, and we built an evaluation around it that addresses the third. Level 1 encodes each report with BiomedBERT [16], a biomedical language model pretrained on PubMed abstracts and PubMed Central full text, adapted with low-rank adaptation (LoRA) [17]. Level 2 is a temporal transformer [12] that reads the sequence of report vectors together with the visit times and conditions on a learnable cancer-type embedding, so that one set of weights can serve many tumour families. Because no fully open corpus contains longitudinal reports for many cancer types, the two capabilities are examined on different data: controlled synthetic benchmarks for temporal modelling, where ground truth and ceilings are known, and the real TCGA-Reports corpus [18,19] for cross-type transfer, where each patient has a single report. We ask four questions.

(a) Does reading the full visit sequence improve prediction over reading the most recent report, and how much improvement is possible?

(b) Does the temporal pathway use the real intervals between visits, and under what conditions can it?

(c) Does tumour-grade prediction transfer to cancer types absent from training, and which component carries the transfer?

(d) How does a text-only HTT compare with a published multimodal survival model on the same cohort?

Our contributions are as follows. First, we describe HTT, a two-level, parameter-efficient architecture for report sequences with type conditioning and a dense per-visit auxiliary loss. Second, we introduce a pair of controlled benchmarks with computable ceilings: a phrase benchmark in which the label is stated in the text and the dates are uninformative, and a growth benchmark in which the label is recoverable only by combining reported sizes with visit intervals. Third, using ablations and re-dating probes, we characterise what the temporal pathway uses, and we show that a time code on a clinical scale fed with the interval since the previous visit (HTT-IA) makes the model respond to visit spacing. Fourth, we evaluate zero-shot transfer on 459 real reports from three withheld tumour types under ten seeds, with paired patient-level uncertainty, and attribute the transfer to the pre-trained encoder. Fifth, we place a text-only HTT against the published multimodal survival model CALM [20] on the same patients and folds. Fig. 1 gives an overview of the study design, and Table 1 situates the work against prior cancer language-model studies.

**Fig. 1.**
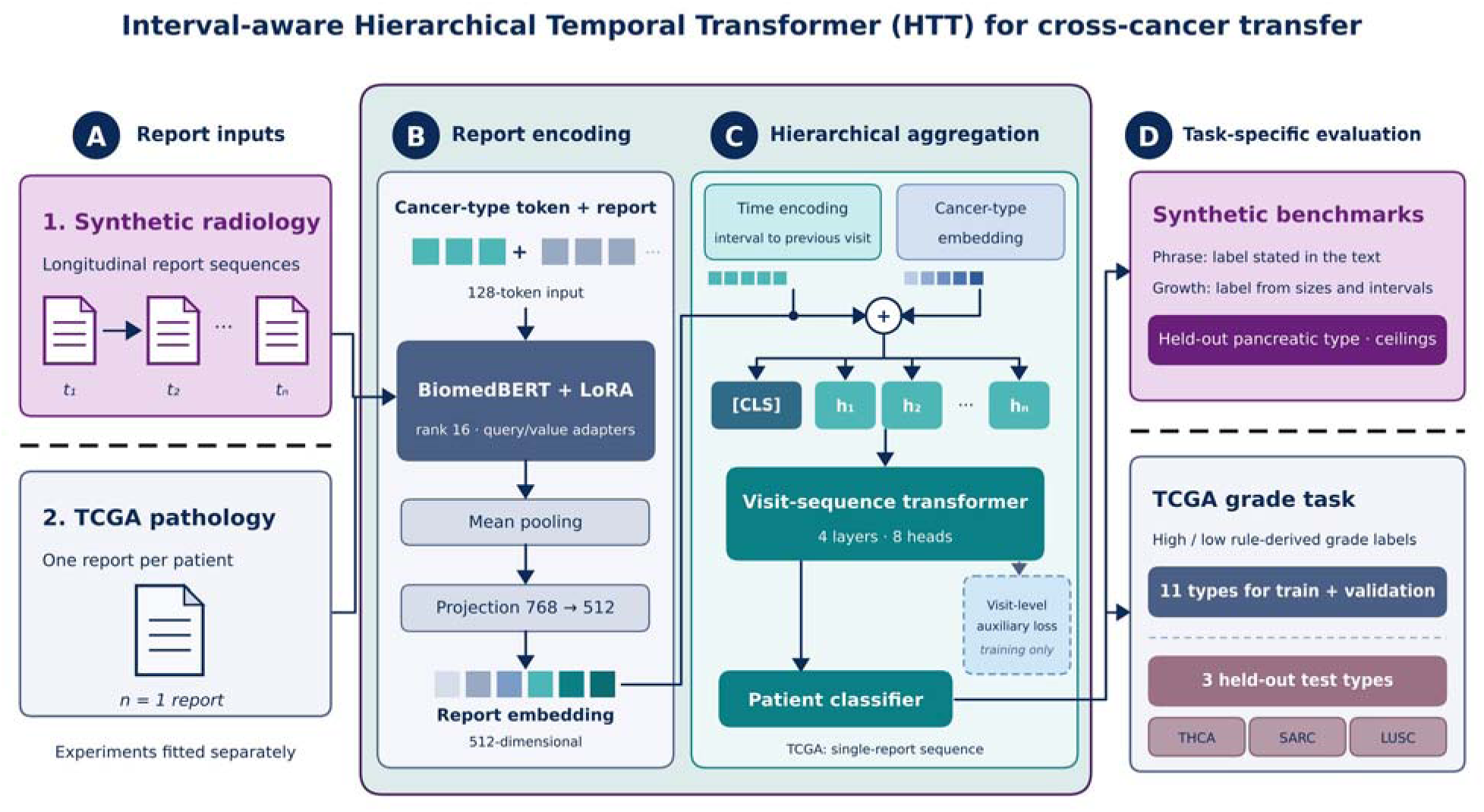
Study overview. (A) Two inputs are fitted separately: synthetic radiology report sequences (phrase and growth benchmarks) and TCGA pathology reports, one per patient. (B) Each report, prefixed with a cancer-type token and cut to 128 tokens, is encoded by BiomedBERT with LoRA, mean-pooled and projected to a 512-dimensional embedding. (C) Report embeddings are combined with a time encoding (elapsed time in the initial model, interval to the previous visit in HTT-IA) and a cancer-type embedding, then read by a visit-sequence transformer with a classification token. A patient classifier gives the prediction, and a visit-level auxiliary loss is used in training only. (D) The synthetic benchmarks hold out one cancer type (pancreatic) and are scored against best-possible AUROC ceilings. The TCGA grade task uses 11 types for training and validation and three held-out types (THCA, SARC, LUSC) for testing; a separate text-only survival comparison with CALM uses 14 types.

**Table 1.** Comparison with prior cancer language-model studies.

| **Study** | **Data** | **Types** | **Temporal** | **Unseen-type evaluation** | **Best AUROC** |
|---|---|---|---|---|---|
| Woollie [3] | Radiology reports | 1 | No | No | 0.80 |
| CancerLLM [4] | Clinical notes | Multiple | No | No | 0.85 |
| Tan et al. [5] | Radiology reports | 1 | No | No | 0.82 |
| Lee et al. [20] | Image and text | Multiple | No | Partial | 0.88 |
| HTT (this work) | Pathology reports; synthetic sequences | 14 (3 held out) | Yes, ceiling-calibrated | Yes | 0.925 (mean, ten seeds) |

AUROC values for prior work are approximate, read from the published figures, and are not directly comparable across tasks.

## 2. Materials and methods

### 2.1 Study design

Fig. 2 shows the models, and Table 2 lists the ten experiments (E0 to E9) that answer the four questions. The experiments share one model family and one training procedure but use different corpora and labels. In every experiment a non-temporal Baseline, which shares the encoder and LoRA configuration and reads only the most recent report, is trained under identical conditions, so that any difference isolates the temporal and conditioning components rather than the pre-trained encoder. All experiments use the source code released with the model (repository rajneeshanand/OncoLocus, commit 47719d1, unmodified); our additions are the growth benchmark, the ceiling calculations, the aggregation and probing scripts, and the time-code variants. Six of the experiments involve training and account for 300 training runs. The parameters of the growth benchmark were fixed after its ceilings were computed and before any model was trained on it, and the protocol of the survival comparison was fixed before any of its results were seen.

**Fig. 2.**
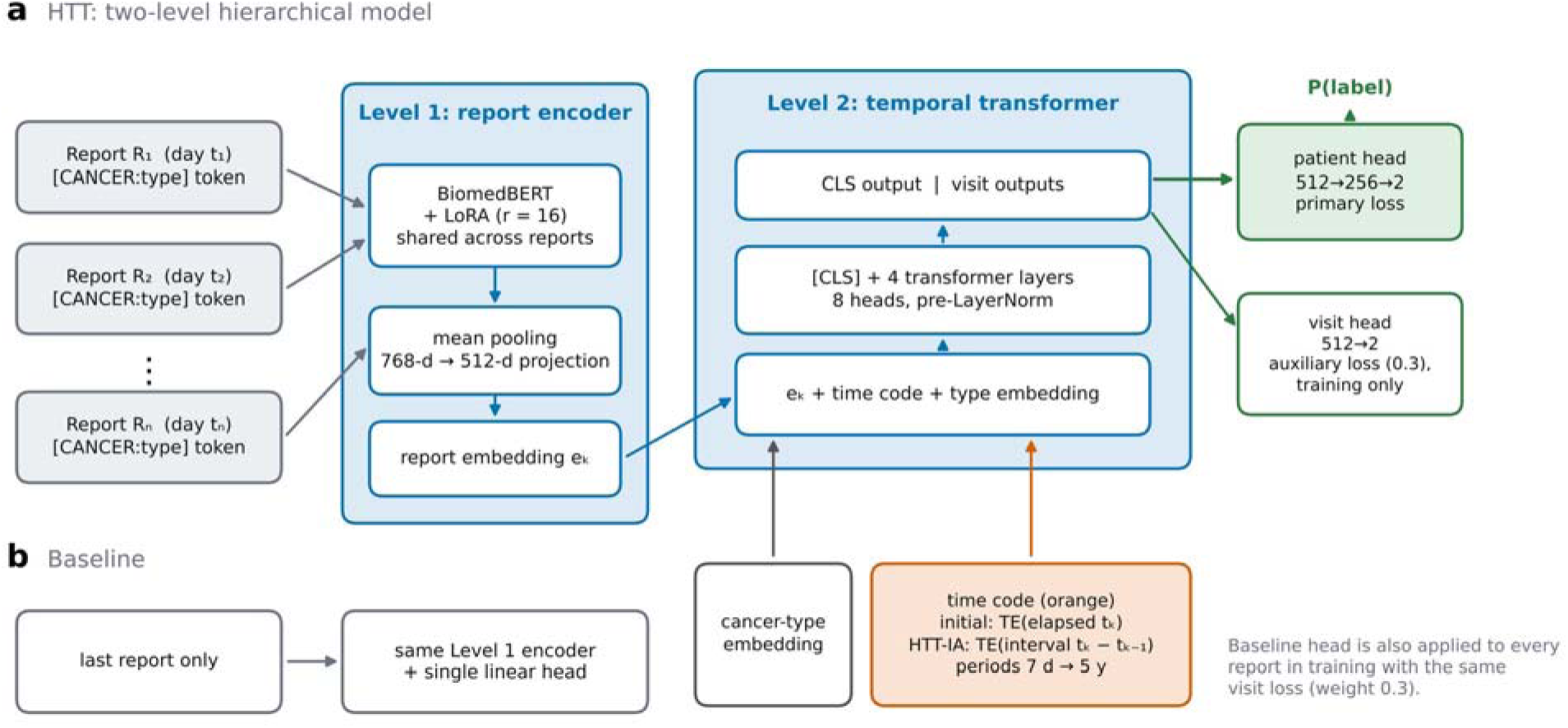
The Hierarchical Temporal Transformer (HTT) and its Baseline. (a) Each report, prefixed with a structured cancer-type token, is encoded independently by BiomedBERT with LoRA, mean-pooled and projected to 512 dimensions (Level 1). The sequence of report embeddings receives a sinusoidal time code, shown in orange, and a learnable cancer-type embedding, and passes with a prepended classification token through a four-layer transformer encoder (Level 2). A patient-level head on the classification output carries the primary loss; a visit-level head on each visit output supplies an auxiliary loss during training only. The initial configuration encodes the elapsed time since the first visit; HTT-IA encodes the interval since the previous visit with periods from 7 days to 5 years. (b) The Baseline reads the most recent report alone through the same Level 1 encoder and a single linear head.

**Table 2.** Experiments.

| ID | Experiment | Question | Data | Runs |
|---|---|---|---|---|
| E0 | Benchmark regeneration | Can the released phrase benchmark be rebuilt and its properties verified? | Phrase benchmark, 500 patients | – |
| E1 | Phrase benchmark | Does HTT beat the last-report baseline with larger test sets and ten seeds? | Phrase; test60/120/180, PANC100/300 | 20 |
| E2 | Temporal ablation | Does HTT use visit order or real intervals? What does the visit loss do? | Phrase | 80 |
| E3 | Re-dating probe | Do predictions change when the same reports are re-dated? | Phrase, test180 | reuse E2 |
| E4 | Ceilings | What is the best AUROC any reader could reach? | Phrase (exact), growth (quadrature) | analytic |
| E5 | Growth benchmark | Can data be built in which timing decides the label, with known ceilings? | Growth | – |
| E6 | HTT on growth benchmark | Does the initial HTT use the dates when they matter? | Growth | 80 |
| E7 | Time-code variants (HTT-IA) | Does a clinical-scale code, or the previous-visit interval, make HTT use the dates? | Growth | 30 |
| E8 | Held-out tumour types | Does grade transfer to unseen types, and does HTT beat its baseline? | TCGA-Reports | 20 |
| E9 | Text-only HTT vs CALM | How does a text-only HTT compare with published multimodal survival results? | TCGA, 14 types, 5 folds | 70 |

Runs are training runs (HTT and Baseline, seeds × conditions). E3 reuses the models of E2; E4 is computed analytically.

### 2.2 Model architecture

HTT comprises a Level 1 report encoder, a Level 2 temporal transformer and two classification heads, and is implemented in PyTorch with the Hugging Face Transformers and PEFT libraries.

Level 1. Each report is encoded independently by BiomedBERT-base, a 110-million-parameter model pretrained from scratch on PubMed abstracts and PubMed Central full text [16]. It was chosen for its biomedical pretraining and open availability and because larger models exceeded the 8 GB of video memory available. LoRA [17] adapts the query and value projections of every attention layer with rank 16, scaling 32 and dropout 0.1, leaving roughly 11.1% of parameters trainable. The last hidden state is mean-pooled over non-padding tokens into a 768-dimensional report embedding and projected to 512 dimensions through a linear layer, layer normalisation and a GELU activation. At inference all reports in a patient’s sequence are encoded in parallel.

Level 2. Three operations are applied to the 512-dimensional report embeddings. First, a continuous-time encoding of the visit time is added. Sinusoidal functions over a bank of 256 frequency channels produce a 512-dimensional vector per visit,

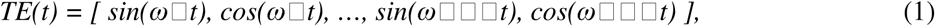

where each frequency carries a learnable scale initialised at 1. In the initial configuration t is the time since the first visit in years and the base frequencies decrease from 1 to 0.05 rad per year, corresponding to periods between 6.3 and 126 years. Second, a learnable 512-dimensional embedding of the patient’s cancer type, with one reserved index for types absent from training, is added to every visit embedding, so that one set of transformer weights can behave differently for a breast, kidney or sarcoma patient; for withheld types the prepended text token still supplies type-specific signal at Level 1. Third, a learnable classification token is prepended and the sequence passes through four transformer encoder layers [12] with 8 attention heads, a feed-forward dimension of 2,048 and pre-layer normalisation, with a padding mask on absent visits. The classification output is the patient-level representation and the remaining outputs are the visit-level representations.

Heads and loss. The patient-level head applies dropout (0.2), a 512→256 linear layer, GELU, dropout (0.2) and a 256→2 layer; the visit-level head applies dropout and a 512→2 layer to each visit output and is used only in training. The loss is the patient-level cross-entropy plus 0.3 times the visit-level cross-entropy, with padded positions excluded. The auxiliary weight lets the visit-level signal guide learning without dominating it, and prevents the model from learning only from the final report.

Baseline. The Baseline shares Level 1 exactly, down to the BiomedBERT weights, LoRA configuration and projection layer. It processes only the most recent report of each patient and passes that embedding through dropout and a single 512→2 layer; it has no temporal transformer, time code or type embedding. During training its head is also applied to every report with the same visit loss of weight 0.3, so that it receives the same dense supervision as HTT. This mirrors the single-report design of prior systems [3,4] and matters for the interpretation of every comparison below. Table 3 lists the variants.

**Table 3.** Model variants.

| **Component** | **Baseline** | **HTT (initial time code)** | **HTT-IA** |
|---|---|---|---|
| Report encoder | BiomedBERT + LoRA (shared) | BiomedBERT + LoRA (shared) | BiomedBERT + LoRA (shared) |
| Reports used per patient | Most recent only | All visits in sequence | All visits in sequence |
| Time input | None | Elapsed time since first visit | Interval since previous visit (0 at first visit) |
| Time-code periods | – | 6.3 to 126 years | 7 days to 5 years (log-spaced) |
| Cancer-type conditioning | None | Learnable embedding | Learnable embedding |
| Temporal transformer | None | 4 layers, 8 heads | 4 layers, 8 heads |
| Patient head | Single linear layer | Two-layer MLP | Two-layer MLP |
| Visit loss (weight 0.3) | Yes | Yes | Yes |
| Trainable parameters | 11.1% of total | 11.1% of total (plus Level 2) | 11.1% of total (plus Level 2) |

The 11.1% figure refers to the LoRA-adapted encoder; Level 2 components are trained in full.

### 2.3 The phrase benchmark

No publicly available corpus contains multiple longitudinal radiology or pathology reports per cancer patient without a data-use agreement, so the temporal capability was first studied on a synthetic corpus of sequential radiology reports (the phrase benchmark). It covers five cancer types: 100 patients each of non-small-cell lung (NSCLC), breast, colorectal and prostate cancer for training, and 100 pancreatic cancer patients withheld entirely for held-out evaluation. Each patient has between two and eight visits and one of four latent trajectories that fix the label of every visit (stable throughout, 30%; progression midway, 30%; progression late, 25%; response then progression, 15%; Appendix Table A1a). The patient label is the label of the last visit. At each visit one of four type-specific templates is chosen, matched to the modality used for that cancer (CT chest for NSCLC, contrast-enhanced breast MRI, CT abdomen and pelvis for colorectal and pancreatic cancer, multiparametric prostate MRI). Variable slots such as lesion size, comparison size, nodal status and modality-specific values (SUVmax, PI-RADS) are filled with plausible sampled values. The disease-change slot is filled deterministically from the trajectory, with phrases such as ‘enlarged’, ‘interval increase’ or ‘progressive disease’ at progressing visits and ‘unchanged’, ‘stable in size’ or ‘no significant interval change’ otherwise. A change slot is present in 4 (breast), 3 (lung), 3 (pancreas), 2 (colorectal) and 1 (prostate) of the four templates per type, which limits what any reader can recover for prostate. A structured cancer-type token is prepended to every report.

Visit k takes place on day D□ = k·g□ , where the gap multiplier g is drawn at each visit from a uniform distribution on 42 to 90 days and is independent of every other variable (Fig. 3). Start dates are randomised between 2018 and 2022. Because the label is written into the text and the dates are unrelated to it, the benchmark tests a narrower question than clinical prediction: whether an architecture that reads the whole sequence can integrate trajectory information that a single-report model cannot access, in a setting where ground truth is known exactly. It does not require the model to use timing, and Section 3.1 quantifies this.

**Fig. 3.**
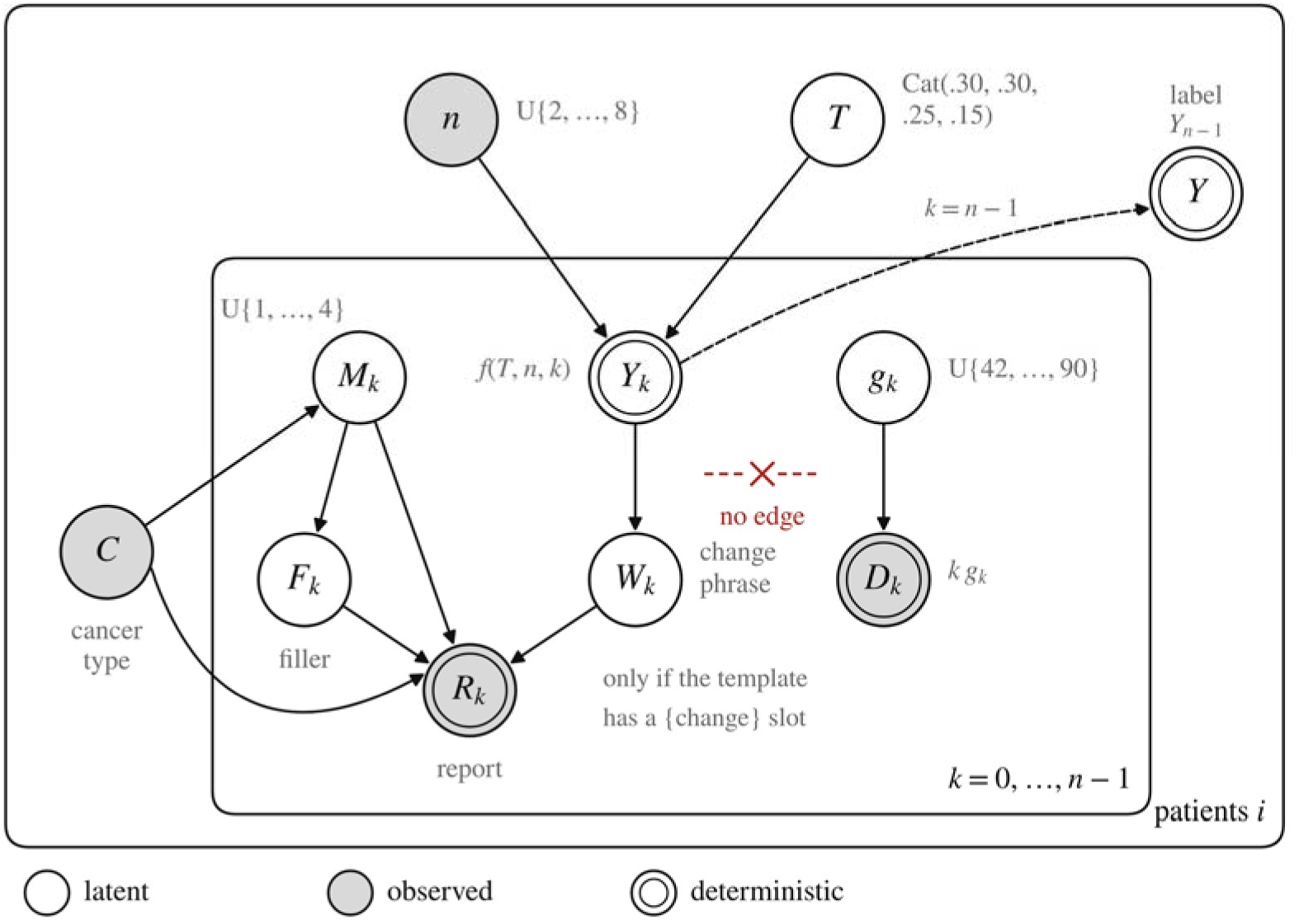
Generative model of the phrase benchmark (plate notation). Shaded circles are what the model receives: cancer type C, visit count n, each report R□ and each visit’s elapsed days D□ . Open circles are latent; double rings are deterministic functions of their parents. The output is the patient label Y, equal to the label of the last visit and never an input. A latent trajectory T and n fix the visit labels Y□ ; Y□ selects the wording W□ of the change phrase, which reaches the report only when the template has a change slot. D□ = k·g□ with g□ independent of everything else, so no path joins the dates to the label (red cross).

Training, validation and test sets follow the original design (Appendix Table A1b): 280 training patients, 60 validation patients, a 60-patient test set (test60) and 100 held-out pancreatic patients (PANC100), with all random operations seeded at 42. To reduce the sampling noise of such small sets we added patients from the same generator under a disjoint seed, giving test120, test180 (our main test set) and PANC300. Training and validation data are identical to the original run.

### 2.4 The growth benchmark

To test the use of timing directly, we built a second generator in which timing decides the label. The label Y is rapid progression (prevalence 0.40). The tumour grows exponentially, s□ = s□·2^(t□ /DT), with a doubling time DT that is log-normal with a median of 120 days when Y = 1 and 600 days otherwise (log-scale standard deviation 0.35). Each patient has a visit cadence between 14 and 180 days and a follow-up limit of 365 days. Each report states the current size and the previous size, each perturbed by 8% multiplicative noise, together with a verdict word derived from their ratio; no date appears in the text (Fig. 4).

**Fig. 4.**
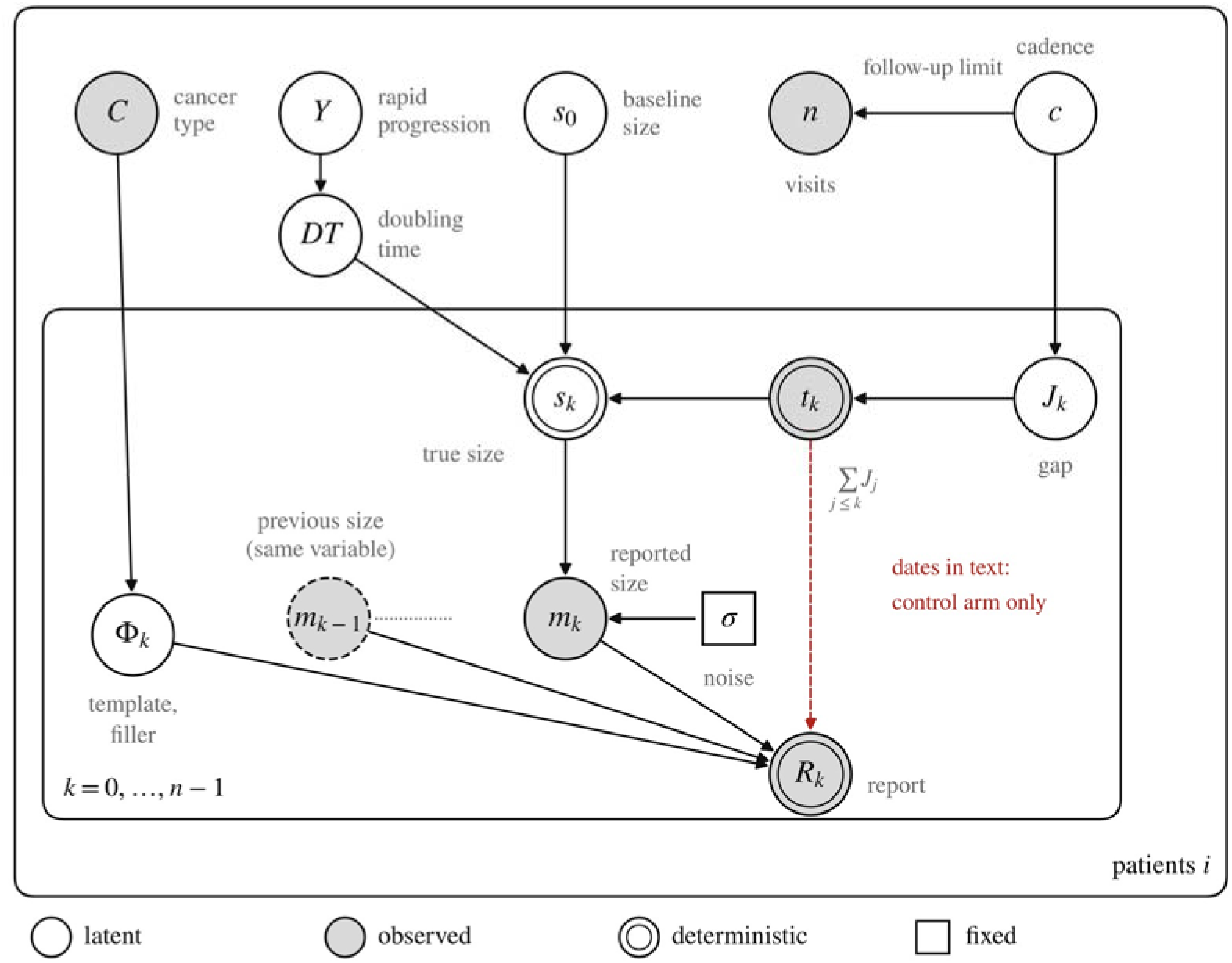
Generative model of the growth benchmark (plate notation). Conventions as in Fig. 3. Shaded circles are inputs: C, n, each report R□ (current size m□ , previous size m□□□ and a verdict) and each date t□ . The label Y sets the doubling time DT; with the baseline size s□ and t□ it fixes the true size s□ , of which the report shows a noisy copy. Dates arise from a patient-level cadence c through the gaps, independently of Y, and the cadence limits n. The dashed red edge (dates printed in the text) is a variant that was not implemented.

Dates are uninformative by themselves, because the cadence is independent of the label, and informative in combination with the sizes. If a report gives sizes s□□□ and s□ and the interval is Δt, then

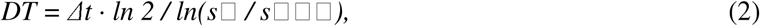

so that the same size ratio implies a short doubling time, and probably Y = 1, over three weeks and a long one over three months. A reader that sees the text but not the dates has to marginalise over the unknown interval; a reader that also sees the dates can apply equation (2). A model therefore has to combine a number read from text with a number supplied as a timestamp. Splits, sizes and construction match the phrase benchmark, and pancreatic cancer is again held out. A preliminary draft with independent gaps rather than a per-patient cadence raised the ceiling by only 0.019 when dates were added, which is why the final generator uses a cadence (Fig. 5b).

**Fig. 5.**
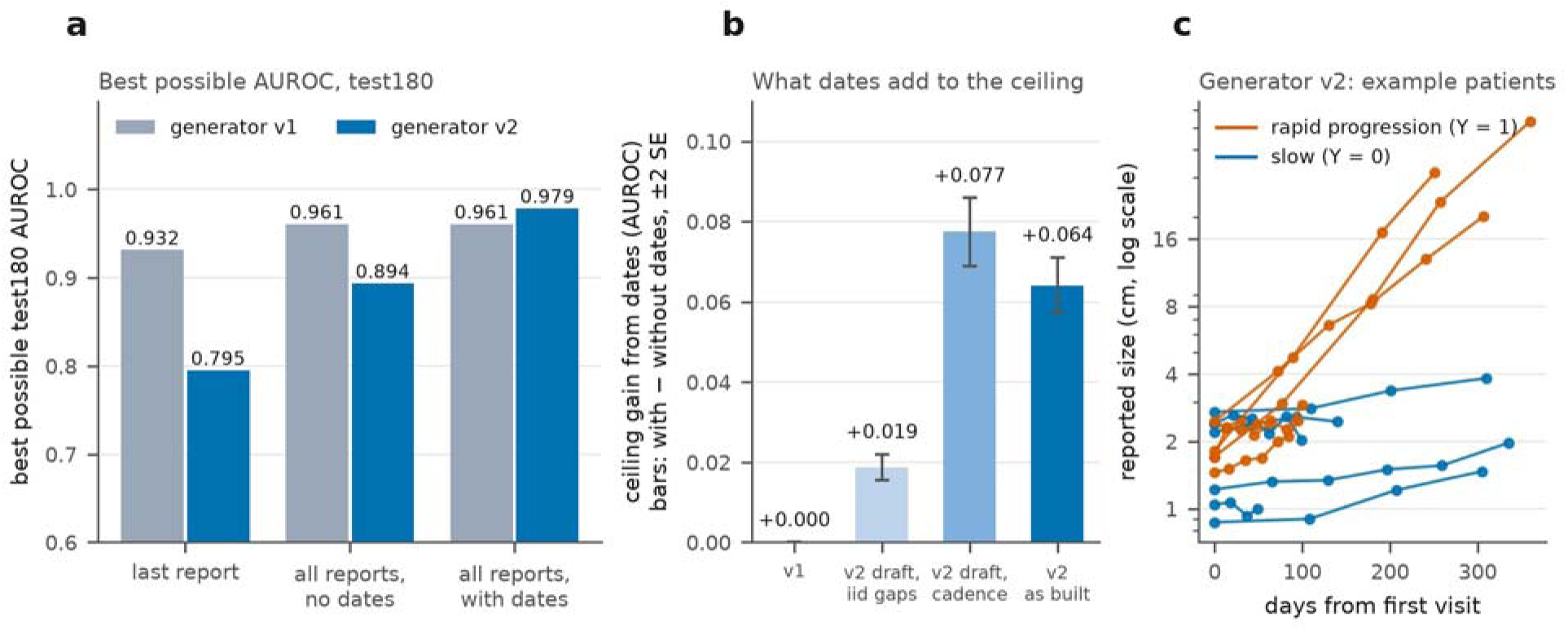
The two benchmarks and what dates can add. (a) Best possible test180 AUROC for the three readers in the two benchmarks (axis starts at 0.6). In the phrase benchmark the dates cannot add anything by construction, so ‘with dates’ equals ‘no dates’. (b) What the dates add to the ceiling (with minus without dates), bars ± 2 standard errors: exactly 0 for the phrase benchmark; two preliminary growth drafts (independent gaps, then a per-patient cadence; 4,000 simulated patients each) and the growth benchmark as built (5,000 simulated patients). (c) Twelve example growth-benchmark patients, reported size on a logarithmic axis: rapid progressors grow faster, but a single size change is ambiguous unless the interval is known.

### 2.5 Ceilings

For a reader with information set I, the best achievable AUROC is that of the true posterior probability π_I(x) = Pr(Y = 1 | I(x)), because AUROC is invariant to monotone transformations and the posterior is monotone in the likelihood ratio [21,22]. We computed this ceiling for three readers: one that sees only the last report, one that sees all reports but no dates, and one that sees all reports and the dates. On the phrase benchmark the posterior can be enumerated exactly over the four trajectories. On the growth benchmark we integrated numerically over the growth rate and sampled the unknown visit gaps for the reader without dates. Ceilings are computed for the specific test patients (sample ceilings) and, for the growth benchmark, for 5,000 simulated patients (population ceiling, with standard error). A sample ceiling is an expectation for a given reader, and on a small sample a model can land above it by chance.

### 2.6 Training protocol, seeds and ablations

Experiments used the configuration of the released code: 128 tokens per report (the longest synthetic report has 59 tokens), AdamW [23] at a learning rate of 2×10 with weight decay 0.01 applied to trainable parameters only, a linear warm-up over the first 10% of steps from 10% of the target rate, batch 4 with 4 accumulation steps, gradient-norm clipping at 1.0, half-precision arithmetic, ten epochs, and early stopping on validation AUROC with a patience of five epochs, restoring the best-validation checkpoint (Appendix Table A2). Validation uses only training types, so no withheld type influences a training decision. Ten training seeds (42 and 1 to 9) were used throughout, and every run was scored on validation, test60, test120, test180, PANC100 and PANC300. Seeds vary training only; the data split is fixed.

Every ablated model remained at chance for several epochs before learning, so a patience of five can end a run before learning has started. We therefore also ran each phrase-benchmark condition for 20 epochs without early stopping and report these separately. The learning take-off is the first epoch at which validation AUROC reaches 0.8. The ablation conditions are the full model; visit loss set to zero; no time (all days set to zero, so that the visits form an unordered set); and order only (days replaced by visit index × 66 days, which preserves order and removes real intervals). Four conditions, ten seeds and two protocols give 80 runs on the phrase benchmark. On the growth benchmark we ran five conditions at ten epochs (full, visit loss zero, no time, order only and the Baseline) and three at 20 epochs (full, no time, Baseline), again with ten seeds, giving 80 runs.

### 2.7 Re-dating probe

For each trained full model we kept the report texts of the 180 test180 patients and re-dated visit k to day k·g for g = 21, 42, 66 and 90 days, and to all days zero, then recorded the predicted probability and the 512-dimensional patient embedding. A model that uses visit pace should change its predictions in a direction consistent with the data-generating process. On the growth benchmark the shift of the ceiling reader between 21 and 90 days (0.374) gives the scale for comparison.

### 2.8 Interval-aware time code (HTT-IA)

The re-dating probe on the phrase benchmark and the learned frequency scales (Section 3.4) indicated that the initial time code behaves as a smooth ramp over a visit history spanning under two years. We therefore replaced it by sinusoids with periods log-spaced from 7 days to 5 years and tried three inputs: the time since the first visit (elapsed), the time since the previous visit, set to 0 at the first visit (interval), and the sum of the two encodings (both). The interval variant is HTT-IA (Table 3). Nothing else changed: the data, protocol, seeds, visit loss and checkpoint rule were the same. One setting of each variant was run for ten epochs. The three variants were fixed in advance and none was tuned on test data, but all were compared on the same test sets.

### 2.9 TCGA tumour-grade experiment

Cross-type transfer was evaluated on TCGA-Reports [18], an open corpus of 9,523 free-text pathology reports from The Cancer Genome Atlas [19] hosted on the Hugging Face Hub without a data-use agreement. Each record pairs a patient identifier with a report covering histology, grade, differentiation, tumour size, margin status and nodal involvement. The corpus holds one report per patient, so the temporal component operates on sequences of length one and this experiment tests the shared encoder and the type conditioning, not temporal modelling.

The corpus has no structured cancer-type labels. We extracted the cancer site from the report text with 17 ordered pattern sets over anatomical and histological keywords, applied most-specific-first so that squamous lung reports are not absorbed by the general lung pattern (representative patterns in Appendix Table A3). Types with at least 40 labelled reports were retained, leaving 14 types. The task is binary: high grade (label 1) against low grade (label 0). High-grade patterns target ‘grade III’, ‘grade 3’, ‘poorly differentiated’, ‘undifferentiated’, ‘high grade’, ‘nuclear grade III’, ‘histologic grade III’ and ‘FNCLCC grade 3’; low-grade patterns target ‘grade I’, ‘grade 1’, ‘well differentiated’, ‘low grade’, ‘histologic grade I’ and ‘FNCLCC grade 1’. A report matching only high-grade patterns is labelled 1 and one matching only low-grade patterns is labelled 0. Reports matching both, usually mixed-grade tumours, are labelled 1, since overall grade is conventionally taken from the highest-grade component, and reports matching neither were excluded. Of 9,523 reports, 4,737 carried no grade keywords, leaving 4,786 labelled reports (3,428 high grade, 1,358 low grade), a mild imbalance that makes AUROC the primary metric. A structured token carrying the project code is prepended to each report.

Splits were made at the patient level. The 11 in-distribution types (BLCA, BRCA, COAD, GBM, HNSC, KIRC, LIHC, LUAD, PRAD, STAD, UCEC) were divided 70/15/15 into training (3,023 reports), validation (644) and test (660). Thyroid carcinoma (THCA), sarcoma (SARC) and lung squamous cell carcinoma (LUSC) were withheld entirely and form a single evaluation set of 459 reports (69, 119 and 271; Appendix Table A4). These three types were chosen because their grading language differs from the training types: thyroid grade is expressed through mitotic activity, necrosis, aggressive variant histology and extrathyroidal extension; sarcoma uses the FNCLCC composite of mitotic count, necrosis and differentiation, a format that appears nowhere in a carcinoma corpus; and lung squamous carcinoma is characterised by keratinisation and intercellular bridges. We trained HTT and the Baseline with ten seeds, saved per-patient predictions, and measured how often the labelling rule finds the same grade phrase within the first 128, 256 and 512 tokens.

### 2.10 Text-only HTT and a multimodal survival model

CALM is a published multimodal model that predicts overall survival from whole-slide image features and LLM-written summaries of the pathology report [20]. We trained a text-only HTT on the same 5,720 patients of the PORPOISE cohort [24] and the same five folds, in 14 cancer types, with CALM’s discrete-time negative log-likelihood, four time bins, validation split and stopping rule (Appendix Table A5). The risk score was the negative summed survival curve and performance was Harrell’s concordance index [25], computed with our own implementation and checked against scikit-survival [26]. The input differed from CALM’s in that we used the raw report text (first 512 tokens) where CALM used LLM-written summaries. CALM numbers are values from its published Table 3 and were not re-run; one seed per fold was used.

### 2.11 Statistical analysis

Differences between models are paired by seed and by patient. Confidence intervals are patient bootstraps [27] with 2,000 resamples of the mean difference over seeds, so that they ask whether a result carries over to new patients; for single TCGA runs we used a class-stratified patient bootstrap. A Wilcoxon signed-rank test across seeds [28] is reported separately; because it treats the evaluation patients as fixed it is the more optimistic of the two, and where it and the bootstrap disagree we say so. AUROC is the primary metric, with F1, precision, recall and accuracy at the default threshold (argmax of the softmax) as secondary metrics, and AUROC values are described with the conventional bands (0.80–0.90 excellent, above 0.90 outstanding) [29]. For the CALM comparison the interval reflects our side only, since the CALM values are fixed. No correction for multiplicity was applied. The software was Python 3.12, torch 2.11, transformers 5.18 and peft 0.21. The phrase-benchmark runs used one RTX A5000 GPU, and the TCGA and growth-benchmark runs a workstation with two RTX 3090 GPUs; a growth-benchmark run takes about 70 seconds.

## 3. Results

### 3.1 Ceilings define what each benchmark can show

Before comparing models we asked what each benchmark allows. On the phrase benchmark the best possible test180 AUROC is 0.932 for a reader that sees only the last report and 0.961 for one that sees all reports, and it is 0.961 again when the dates are added, because given the visit count the dates are independent of the label (Fig. 5a,b). The only information carried by timing is the visit count itself, which carries a little (two-visit patients are positive in 47% of cases, against 61% to 80% for other counts). All reports can therefore raise the ceiling by at most 0.029, and timing cannot raise it at all. In this regime a model that reads the sequence can gain from aggregation, but a model that ignores the dates is behaving optimally.

The growth benchmark removes this limitation. On its test180 set the ceiling is 0.795 for the last report, 0.894 for all reports without dates and 0.979 with dates; in the population the values are 0.833 ± 0.006, 0.922 ± 0.004 and 0.986 ± 0.001 (Table 4). Dates add 0.064 to 0.085 to the ceiling, so a model that reads them can move well beyond one that reads only text. The two benchmarks thus provide a regime in which timing is irrelevant and one in which it is decisive, with known headroom in each.

**Table 4.** Growth benchmark: evaluation sets and best possible AUROC of three readers.

| **Set** | **Patients** | **Positive rate** | **Ceiling: last report** | **Ceiling: all reports, no dates** | **Ceiling: all reports, with dates** |
|---|---|---|---|---|---|
| Validation | 60 | 0.42 | 0.893 | 0.928 | 0.989 |
| test60 | 60 | 0.48 | 0.720 | 0.865 | 0.992 |
| test120 | 120 | 0.43 | 0.804 | 0.904 | 0.987 |
| test180 | 180 | 0.40 | 0.795 | 0.894 | 0.979 |
| PANC100 | 100 | 0.43 | 0.743 | 0.891 | 0.980 |
| PANC300 | 300 | 0.39 | 0.815 | 0.911 | 0.983 |
| **Population (5,000 simulated)** | **5,000** | **0.40** | **0.833 ± 0.006** | **0.922 ± 0.004** | **0.986 ± 0.001** |

Values for the named sets are sample ceilings for those patients; the last row is the population value with its standard error.

### 3.2 On the phrase benchmark, reading the sequence helps modestly

The released benchmark was regenerated exactly: split sizes and class counts match the original confusion matrices, and file hashes are identical on two machines. Its properties agree with the specification of the generator (Appendix Table A6); in particular, visit dates are not cumulative, because the gap multiplier is redrawn at each visit, so that 10% of gaps are zero or negative and 35% of patients have out-of-order dates. Visits are fed to the model in index order. The seed-42 run closely follows the original run (Appendix Table A7): best validation AUROC 0.939 against 0.942, test60 0.927 against 0.912, PANC100 0.998 against 0.995. HTT leads on validation and on the held-out pancreatic patients, and the Baseline is slightly ahead on the 60-patient test set, where a single reclassified patient moves AUROC by about two points.

Over ten seeds the picture is more detailed (Table 5; Fig. 6). With 180 test patients HTT reaches 0.931 ± 0.012 against 0.913 ± 0.007, a paired difference of +0.018 (95% CI −0.009 to +0.050) with HTT ahead in 9 of 10 seeds. The Wilcoxon test across seeds gives p = 0.014, but the patient bootstrap includes zero. On test60 and test120 the sign reverses (−0.012 and −0.006). On the held-out pancreatic patients HTT leads (PANC100 +0.032, 95% CI +0.002 to +0.070; PANC300 +0.019, −0.004 to +0.042), in line with the expectation that a single-report model has least to go on for a type it has not seen.

**Table 5.** Phrase benchmark: test AUROC over ten training seeds, and HTT minus Baseline.

| **Set** | **n** | **HTT, mean ± SD** | **Baseline, mean ± SD** | **HTT − baseline [95% CI]** | **Seeds HTT > base** | **Wilcoxon p** | **Room for gain** |
|---|---|---|---|---|---|---|---|
| test60 | 60 | 0.904 ± 0.025 | 0.916 ± 0.015 | −0.012 [−0.068, +0.044] | 4/10 | 0.36 | 0.039 |
| test120 | 120 | 0.922 ± 0.018 | 0.928 ± 0.008 | −0.006 [−0.041, +0.026] | 5/10 | 0.56 | 0.021 |
| test180 | 180 | 0.931 ± 0.012 | 0.913 ± 0.007 | +0.018 [−0.009, +0.050] | 9/10 | 0.014 | 0.029 |
| Validation | 60 | – | – | +0.062 [+0.007, +0.132] | 10/10 | 0.002 | 0.066 |
| PANC100 | 100 | 0.984 ± 0.019 | 0.952 ± 0.016 | +0.032 [+0.002, +0.070] | 9/10 | 0.004 | 0.029 |
| PANC300 | 300 | 0.955 ± 0.017 | 0.936 ± 0.012 | +0.019 [−0.004, +0.042] | 9/10 | 0.027 | 0.024 |

Differences are paired by seed and patient; intervals are patient bootstraps (2,000 resamples) of the mean difference over seeds. Room for gain is the all-reports ceiling minus the last-report ceiling. Validation is the checkpoint-selection set.

**Fig. 6.**
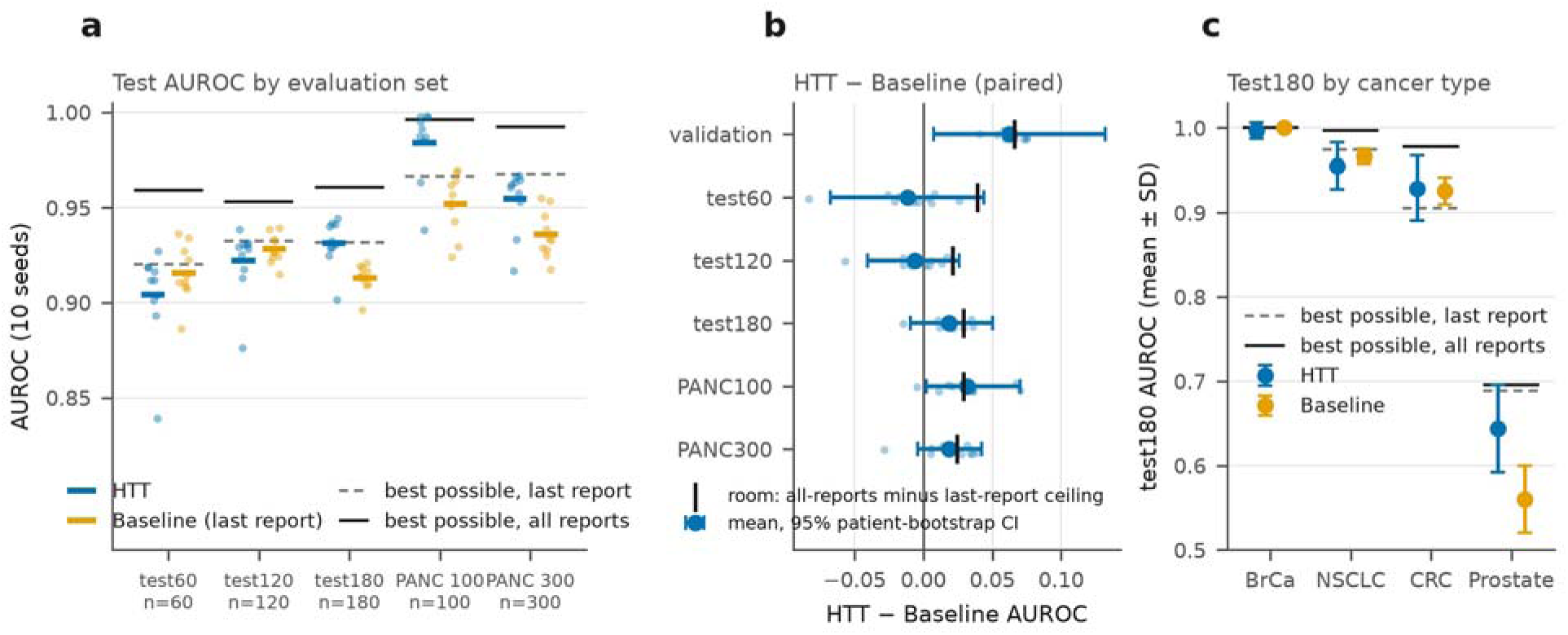
Phrase benchmark: HTT and the last-report Baseline across ten seeds. (a) Test AUROC; dots are seeds and bars the means; grey dashed and black lines are the best possible AUROC from the last report and from all reports for those patients. (b) HTT minus Baseline: dots per seed, circle the mean, bars the 95% patient-bootstrap CI, vertical tick the room for gain. (c) test180 by cancer type, mean ± SD over seeds, with the per-type ceilings.

Two features of the data explain much of this. Half of the validation patients have an uninformative last report, against 38% in test180, so HTT has more to gain there (+0.062 of the 0.066 available); validation is also the checkpoint-selection set and is not independent evidence. And the room for gain is small to begin with: the difference between the all-reports and last-report ceilings is 0.029 on test180, of the same order as the sampling noise of a small test set (the width of a single-run 95% interval falls from 0.137 at 60 patients to 0.068 at 180). PANC100 was an easy draw (0.984), which falls to 0.955 when the set is tripled. Per-type results (Appendix Table A8) show breast cancer at ceiling (0.997), NSCLC at 0.956 and colorectal at 0.929, and prostate near 0.64, below its ceiling of 0.69, because only one of its four templates carries a change phrase.

### 3.3 Visit order speeds learning, and the visit loss is decisive

Table 6 and Fig. 7 show the ablation. Removing the dates changes the final result by nothing measurable. With 20 epochs, test180 AUROC is 0.936 without any time input against 0.937 for the full model (−0.001, 95% CI −0.009 to +0.007), and order alone gives 0.931 (−0.006, −0.013 to +0.000). Order does speed early learning: at epoch 7 the validation AUROC is 0.840 without time and 0.926 with the full code, lower in 10 of 10 seeds (Wilcoxon p = 0.002), and the order-only model matches the full model (0.926). This is what the ceilings predict for a benchmark in which the dates carry no information beyond the visit sequence.

**Table 6.** Phrase benchmark: temporal ablation on test180 under two training protocols.

| **Condition** | **Published protocol, test180** | **Seeds that learned** | **20 epochs, test180** | **Δ vs full [95% CI]** | **Take-off epoch (range)** |
|---|---|---|---|---|---|
| Full | 0.931 ± 0.012 | 10/10 | 0.937 ± 0.009 | – | 6.8 (6–7) |
| Visit loss = 0 | 0.558 ± 0.058 | 0/10 | 0.924 ± 0.019 | −0.013 [−0.023, −0.004] | 12.6 (11–14) |
| No time | 0.890 ± 0.130 | 9/10 | 0.936 ± 0.005 | −0.001 [−0.009, +0.007] | 7.2 (7–8) |
| Order only | 0.931 ± 0.009 | 10/10 | 0.931 ± 0.016 | −0.006 [−0.013, +0.000] | 6.8 (6–7) |
| Baseline | 0.913 ± 0.007 | 10/10 | – | – | 5.1 (5–6) |

Mean ± SD over ten seeds. Δ vs full is the paired difference of the 20-epoch runs with a 95% patient-bootstrap CI. Take-off is the first epoch with validation AUROC ≥ 0.8. ‘Seeds that learned’ refers to the published protocol (10 epochs, patience 5).

**Fig. 7.**
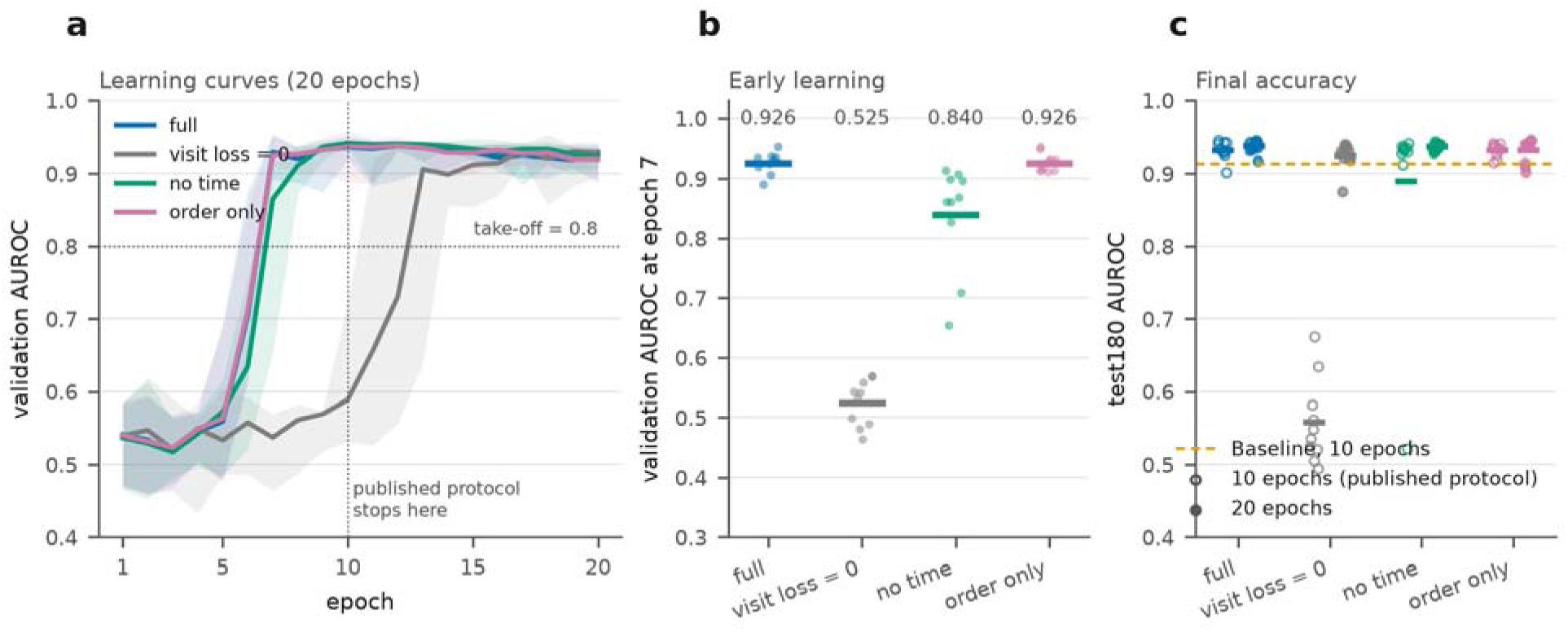
Phrase benchmark: temporal ablation. (a) Validation AUROC per epoch with 20 epochs and no early stopping: median line and min–max band over ten seeds; dotted lines mark take-off (0.8) and the point at which the published protocol stops (epoch 10). (b) Validation AUROC at epoch 7 for each seed (numbers are means). (c) test180 AUROC under the published protocol (open circles) and with 20 epochs (filled); the dashed line is the Baseline’s ten-epoch mean.

The visit loss matters more than the time code. Under the published protocol the model without it never learned in any of ten seeds (test180 AUROC 0.558), although every seed learned when given 20 epochs, six epochs later and 0.013 AUROC lower (0.924 against 0.937; −0.013, 95% CI −0.023 to −0.004). The loss provides about 1,400 per-report labels per epoch in place of 280 patient labels, and its absence turns a delay in learning into a failed run, because early stopping with a patience of five ends training during the initial plateau.

The same plateau explains the training dynamics. Validation AUROC of the full model hovers between 0.42 and 0.66 for six epochs and then jumps at epoch 7. The jump is not specific to the temporal pathway: it also occurs when time is removed (take-off at epoch 7.2) and in the Baseline (epoch 5.1). It reflects how the encoder and head leave an initial plateau under a dense per-visit loss, and it should not be read as a signature of temporal reasoning.

### 3.4 The initial time code represents visit pace but barely uses it

We re-dated the reports of the 180 test patients and recorded the predictions of the ten full models (Table 7; Fig. 8). Predictions hardly moved: the mean absolute change between 21-day and 90-day spacing was 0.039 ± 0.019, class labels flipped for 3.9% of predictions, and AUROC was 0.928, 0.931, 0.933 and 0.933 at 21, 42, 66 and 90 days. The change was in the wrong direction for a rate signal. Faster visits lowered the predicted probability of progression for everyone (−0.039 in progressors and −0.038 in non-progressors), because a longer elapsed time looks like ‘more visits’ and patients with more visits are slightly more often positive.

**Table 7.** Phrase benchmark: change in prediction when the same reports are re-dated.

| **Metric (ten full models)** | **Value** |
|---|---|
| Mean \|P(21 d) − P(90 d)\| | 0.039 ± 0.019 |
| Mean P(21 d) − P(90 d), progressors / non-progressors | −0.039 / −0.038 |
| Predictions changing class, 21 vs 90 d | 3.9% |
| test180 AUROC at 21 / 42 / 66 / 90 days | 0.928 / 0.931 / 0.933 / 0.933 |
| test180 AUROC with all days = 0 | 0.921 |
| 21 vs 90 d decodable from the patient embedding | 93% (chance 50%) |
| Pace shift ÷ progressor/non-progressor gap | 0.09 |

Published-protocol full models on test180 patients; the text of each report is unchanged.

**Fig. 8.**
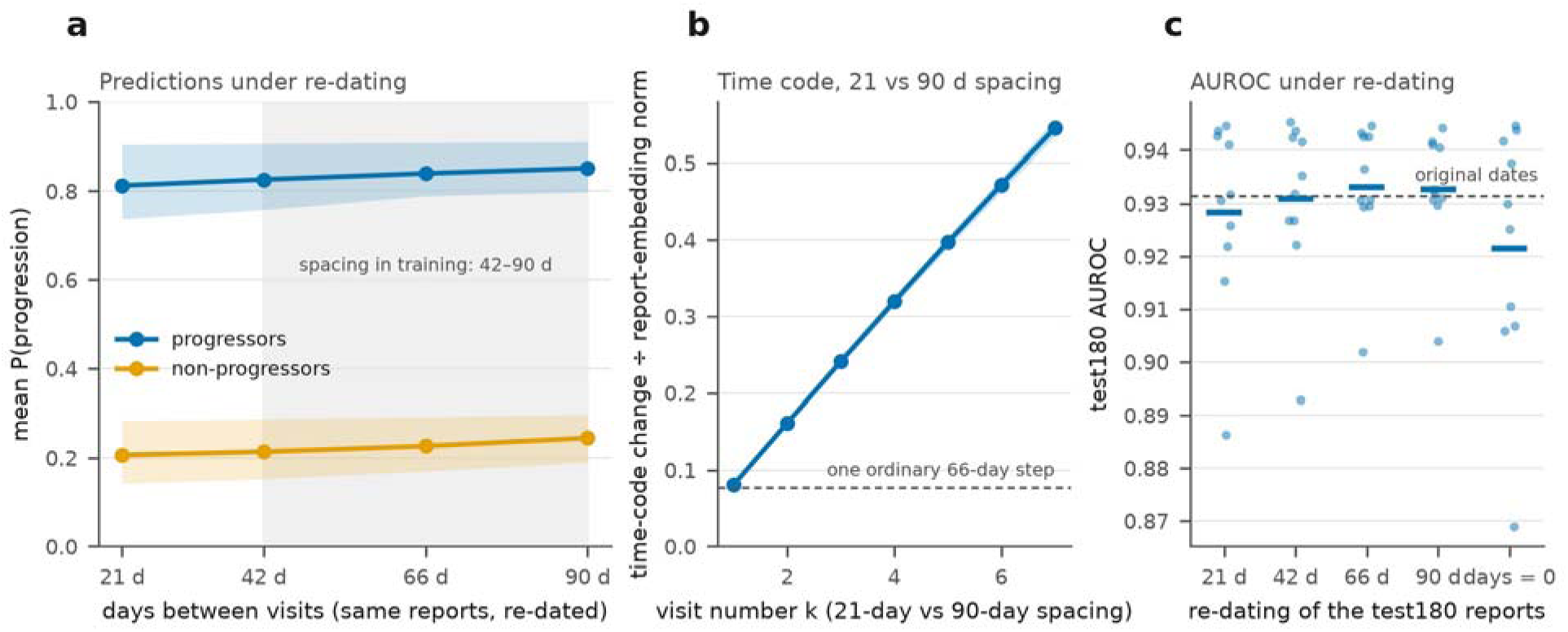
Phrase benchmark: re-dating probe. (a) Mean predicted probability of progression for progressors and non-progressors when the reports are re-dated every 21 to 90 days; bands are the range over ten models; grey marks the spacing range of the training data. (b) Size of the change in the time code between 21-day and 90-day spacing, relative to the report-embedding norm, by visit number (band: SD); the dashed line is one ordinary 66-day step. (c) test180 AUROC under each re-dating; dots are models.

The pace is nevertheless represented. A 21-day spacing can be told from a 90-day spacing from the patient embedding 93% of the time (chance 50%), along one ordered, nearly linear direction (Appendix Table A9). That direction is about one tenth the size of the progressor/non-progressor gap, so it never decides a prediction. The time code had also stayed close to its initialisation: after training the learnable frequency scales averaged 1.004. Since the base frequencies correspond to periods of 6.3 to 126 years while visits span under two years, the code behaves like a smooth ramp in elapsed time and cannot separate three weeks from three months at the first follow-up. This motivated the interval-aware code of Section 2.8.

### 3.5 When timing decides the label, the initial code is not used and the interval-aware code is

We ran the phrase-benchmark conditions on the growth benchmark under the published protocol with ten seeds each (Table 8; Fig. 9). HTT with the initial time code reached a test180 AUROC of 0.790 ± 0.017. Without any time input the score was 0.787 ± 0.011 (−0.003, 95% CI −0.009 to +0.004) and with order only 0.792. All three lie at the last-report ceiling of 0.795 and far below the 0.894 attainable without dates and the 0.979 attainable with them. Re-dating the reports changed the predictions of HTT by 0.010 between 21-day and 90-day spacing, against 0.374 for the best possible reader, and the learned frequency scales again stayed close to 1. This is not an effect of training length: with 20 epochs HTT scored 0.803 and the no-time model 0.798 (−0.006, −0.013 to +0.001), and the Baseline 0.775 (Appendix Table A10). Against the Baseline, HTT is ahead on every set (test180 +0.010, 95% CI −0.041 to +0.059, ahead in 5 of 10 seeds), but every patient-bootstrap interval includes zero. The visit loss helps again: without it test180 AUROC is 0.760 (−0.030, −0.041 to −0.020), and learning takes off later (7.9 against 6.4 epochs).

**Table 8.** Growth benchmark: AUROC of all models on all sets (published protocol, ten seeds).

| **Model** | **Validation** | **test60** | **test120** | **test180** | **PANC100** | **PANC300** |
|---|---|---|---|---|---|---|
| Baseline (last report) | 0.874 ± 0.005 | 0.727 ± 0.025 | 0.794 ± 0.016 | 0.780 ± 0.017 | 0.771 ± 0.012 | 0.826 ± 0.009 |
| HTT, visit loss = 0 | 0.875 ± 0.015 | 0.745 ± 0.039 | 0.792 ± 0.042 | 0.760 ± 0.039 | 0.794 ± 0.025 | 0.829 ± 0.016 |
| HTT, no time input | 0.888 ± 0.007 | 0.769 ± 0.017 | 0.818 ± 0.013 | 0.787 ± 0.011 | 0.795 ± 0.028 | 0.831 ± 0.019 |
| HTT, order only | 0.886 ± 0.012 | 0.773 ± 0.017 | 0.820 ± 0.016 | 0.792 ± 0.011 | 0.799 ± 0.029 | 0.836 ± 0.020 |
| HTT, original time code | 0.890 ± 0.010 | 0.772 ± 0.013 | 0.817 ± 0.020 | 0.790 ± 0.017 | 0.814 ± 0.022 | 0.845 ± 0.018 |
| HTT, clinical code: elapsed | 0.899 ± 0.014 | 0.813 ± 0.026 | 0.839 ± 0.029 | 0.810 ± 0.021 | 0.800 ± 0.035 | 0.838 ± 0.027 |
| HTT, clinical code: elapsed + interval | 0.898 ± 0.020 | 0.862 ± 0.049 | 0.876 ± 0.041 | 0.830 ± 0.044 | 0.845 ± 0.031 | 0.873 ± 0.027 |
| HTT, clinical code: interval | 0.917 ± 0.012 | 0.868 ± 0.034 | 0.880 ± 0.021 | 0.843 ± 0.026 | 0.846 ± 0.029 | 0.877 ± 0.021 |
| **Best possible, last report** | **0.893** | **0.720** | **0.804** | **0.795** | **0.743** | **0.815** |
| **Best possible, no dates** | **0.928** | **0.865** | **0.904** | **0.894** | **0.891** | **0.911** |
| **Best possible, with dates** | **0.989** | **0.992** | **0.987** | **0.979** | **0.980** | **0.983** |

Mean ± SD over ten seeds. ‘Clinical code’ denotes periods log-spaced from 7 days to 5 years; elapsed, interval and elapsed + interval refer to the time input. Ceilings are sample values.

**Fig. 9.**
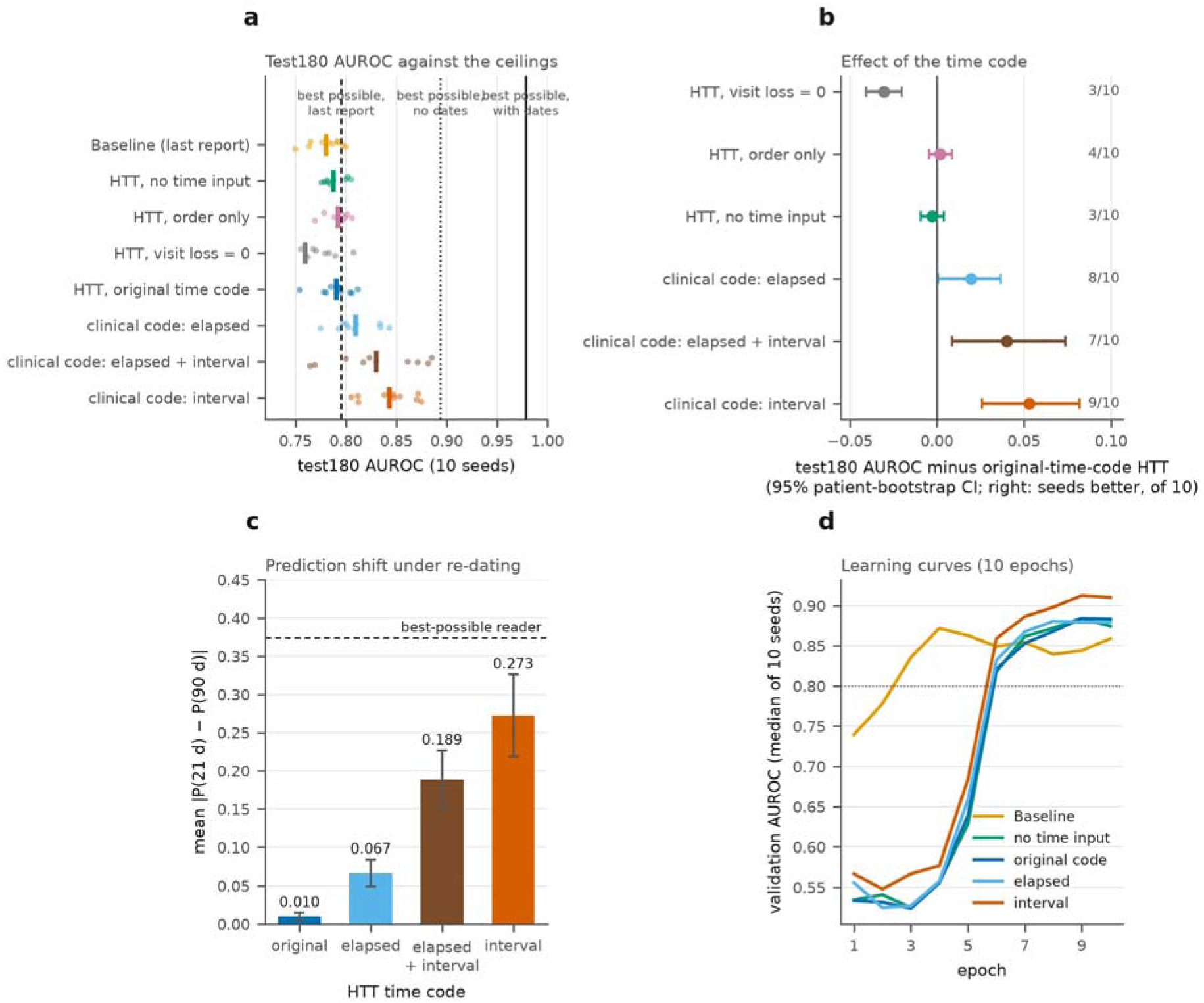
Growth benchmark: does HTT use the visit dates? (a) test180 AUROC for each model (dots: seeds; bar: mean) against the best possible AUROC with the last report, without dates and with dates. (b) Difference from the initial-time-code HTT with 95% patient-bootstrap CIs; numbers on the right give seeds better, of ten. (c) Mean absolute change in predicted probability when the test180 reports are re-dated every 21 against every 90 days (bars: mean over models; whiskers: SD); the dashed line is the best possible reader’s change. (d) Validation AUROC per epoch (median over ten seeds). ‘Clinical code’ denotes periods from 7 days to 5 years; ‘interval’ is HTT-IA.

HTT-IA changes this (Table 9; Fig. 9). The interval-only variant reached a test180 AUROC of 0.843 ± 0.026 (+0.053 over the initial HTT, 95% CI +0.026 to +0.082, better in 9 of 10 seeds; +0.063 over the Baseline, +0.011 to +0.117). Its predictions moved by 0.273 between 21-day and 90-day spacing, about three quarters of the shift of the best possible reader (Fig. 9c), and validation AUROC moved in the same way (0.890 to 0.917). Elapsed time alone with the clinical-scale periods helped little (0.810, +0.020, 95% CI +0.001 to +0.037; shift 0.067), and combining both inputs gave 0.830 but was the least stable, with seeds spanning 0.764 to 0.885.

**Table 9.** Growth benchmark, test180: effect of the time code.

| Model | test180 AUROC, mean ± SD (range) | Δ vs original HTT [95% CI]; seeds better | Δ vs baseline [95% CI] | Take-off epoch | Prediction shift, 21 vs 90 d |
|---|---|---|---|---|---|
| HTT, original time code | 0.790 ± 0.017 (0.754–0.812) | – | +0.010 [−0.042, +0.058] | 6.4 | 0.010 |
| HTT, no time input | 0.787 ± 0.011 (0.775–0.805) | −0.003 [−0.009, +0.004]; 3/10 | +0.007 [−0.042, +0.057] | 6.3 | – |
| HTT, clinical code: elapsed | 0.810 ± 0.021 (0.775–0.843) | +0.020 [+0.001, +0.037]; 8/10 | +0.029 [−0.023, +0.082] | 5.9 | 0.067 |
| HTT, clinical code: interval | 0.843 ± 0.026 (0.806–0.875) | +0.053 [+0.026, +0.082]; 9/10 | +0.063 [+0.011, +0.117] | 5.8 | 0.273 |
| HTT, clinical code: elapsed + interval | 0.830 ± 0.044 (0.764–0.885) | +0.040 [+0.009, +0.074]; 7/10 | +0.050 [+0.001, +0.101] | 5.6 | 0.189 |
| Baseline (last report) | 0.780 ± 0.017 (0.749–0.799) | – | – | 2.5 | – |

Ten seeds each; intervals are patient bootstraps of the mean paired difference. Prediction shift is the mean absolute change in P(progression) when the same reports are re-dated every 21 against every 90 days; the best possible reader’s shift is 0.374. Ceilings on test180: 0.894 without dates, 0.979 with dates.

The improvement is partial. HTT-IA remains 0.051 below the ceiling without dates and 0.136 below the ceiling with dates, so the model sees the dates but extracts only part of what they offer. The interval-only variant has no access to visit position or elapsed time and still performs best, so ordering information was not needed on this benchmark. We did not run the original frequencies with the interval input, so we cannot say how much of the gain comes from the input and how much from the new frequency range. All variants were run for ten epochs only, and the best of three variants on test180 carries some selection optimism, although the validation set shows the same ordering.

### 3.6 Real reports: grade is predicted for tumour types never seen in training

On the 660-report in-distribution test set spanning 11 types, a single set of weights with no per-type head and no per-type fine-tuning reached AUROC 0.923 in the seed-42 run (F1 0.905, precision 0.881, recall 0.929, accuracy 0.853, specificity 0.624); over ten seeds the in-distribution AUROC was 0.928 ± 0.006. In other words, the model ranks a random high-grade report above a random low-grade one about 92% of the time across a biologically diverse set of tumours. Specificity is the weak point: the corpus runs about 2.5:1 high to low grade and the model has absorbed that prior, catching 93% of high-grade cases while correctly identifying 62% of low-grade ones. Per-type AUROC in the seed-42 run spans a wide band (Table 10): kidney clear cell carcinoma leads at 0.971, followed by bladder and colon at 0.951 each, and lung adenocarcinoma is the weakest at 0.710. Prostate is undefined because every prostate report in the test split was high grade.

**Table 10.** TCGA tumour grade: HTT per-type performance on the in-distribution test set (seed-42 run).

| **Code** | **Cancer type** | **AUROC** | **F1** | **n** |
|---|---|---|---|---|
| KIRC | Kidney renal clear cell carcinoma | 0.971 | 0.941 | 85 |
| BLCA | Bladder urothelial carcinoma | 0.951 | 0.989 | 94 |
| COAD | Colon adenocarcinoma | 0.951 | 0.919 | 47 |
| UCEC | Uterine corpus endometrial carcinoma | 0.912 | 0.904 | 92 |
| LIHC | Liver hepatocellular carcinoma | 0.903 | 0.700 | 28 |
| STAD | Stomach adenocarcinoma | 0.894 | 0.935 | 36 |
| BRCA | Breast invasive carcinoma | 0.873 | 0.884 | 87 |
| HNSC | Head and neck squamous cell carcinoma | 0.873 | 0.816 | 39 |
| GBM | Glioblastoma multiforme | 0.846 | 0.804 | 87 |
| LUAD | Lung adenocarcinoma | 0.710 | 0.889 | 37 |
| PRAD | Prostate adenocarcinoma | undefined | 1.000 | 28 |

AUROC is undefined for PRAD because all prostate test samples were high grade; it is excluded from averages. Overall in-distribution AUROC 0.923 (n = 660).

The central result concerns the three withheld types. No report from thyroid carcinoma, sarcoma or lung squamous carcinoma was used in training, validation or hyperparameter selection. Over ten seeds the macro-average AUROC was 0.925 ± 0.007 (published single run 0.923), with a patient-bootstrap interval of 0.894 to 0.954, matching the in-distribution value of 0.928 ± 0.006 (Table 11; Fig. 10a): there is no measurable penalty for transfer at these sample sizes. By the conventional bands two types are outstanding and the third excellent: thyroid 0.996 ± 0.010, sarcoma 0.960 ± 0.014 and lung squamous carcinoma 0.819 ± 0.018. The uncertainty lies in the patients and not in the seeds, as the interval for lung squamous carcinoma (0.737 to 0.899) shows, and thyroid carcinoma has only eight positive cases, so its interval collapses at an AUROC of 1.000 and is uninformative.

**Table 11.**
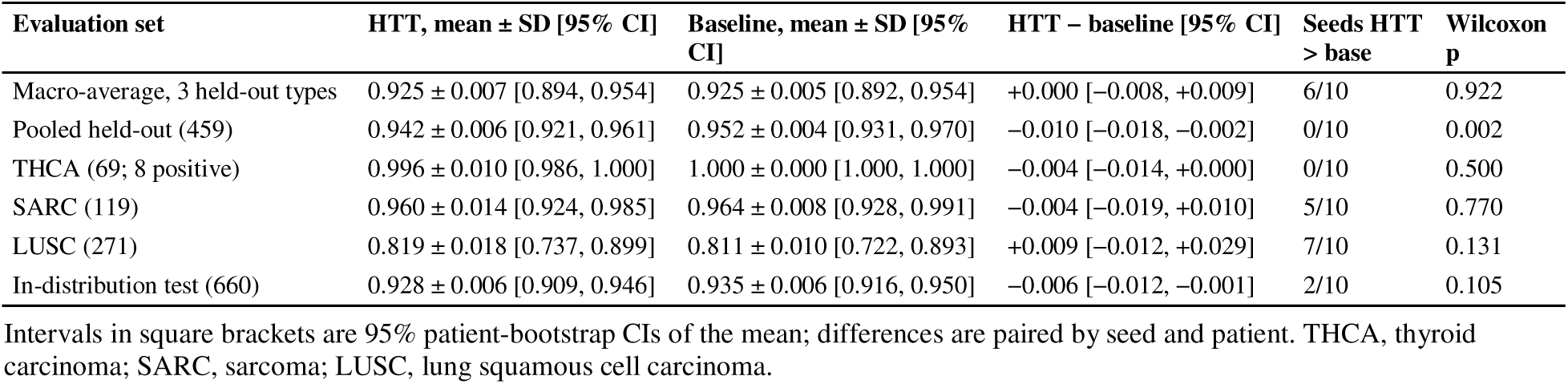
TCGA tumour grade: held-out AUROC over ten seeds, HTT against the Baseline.

**Fig. 10.**
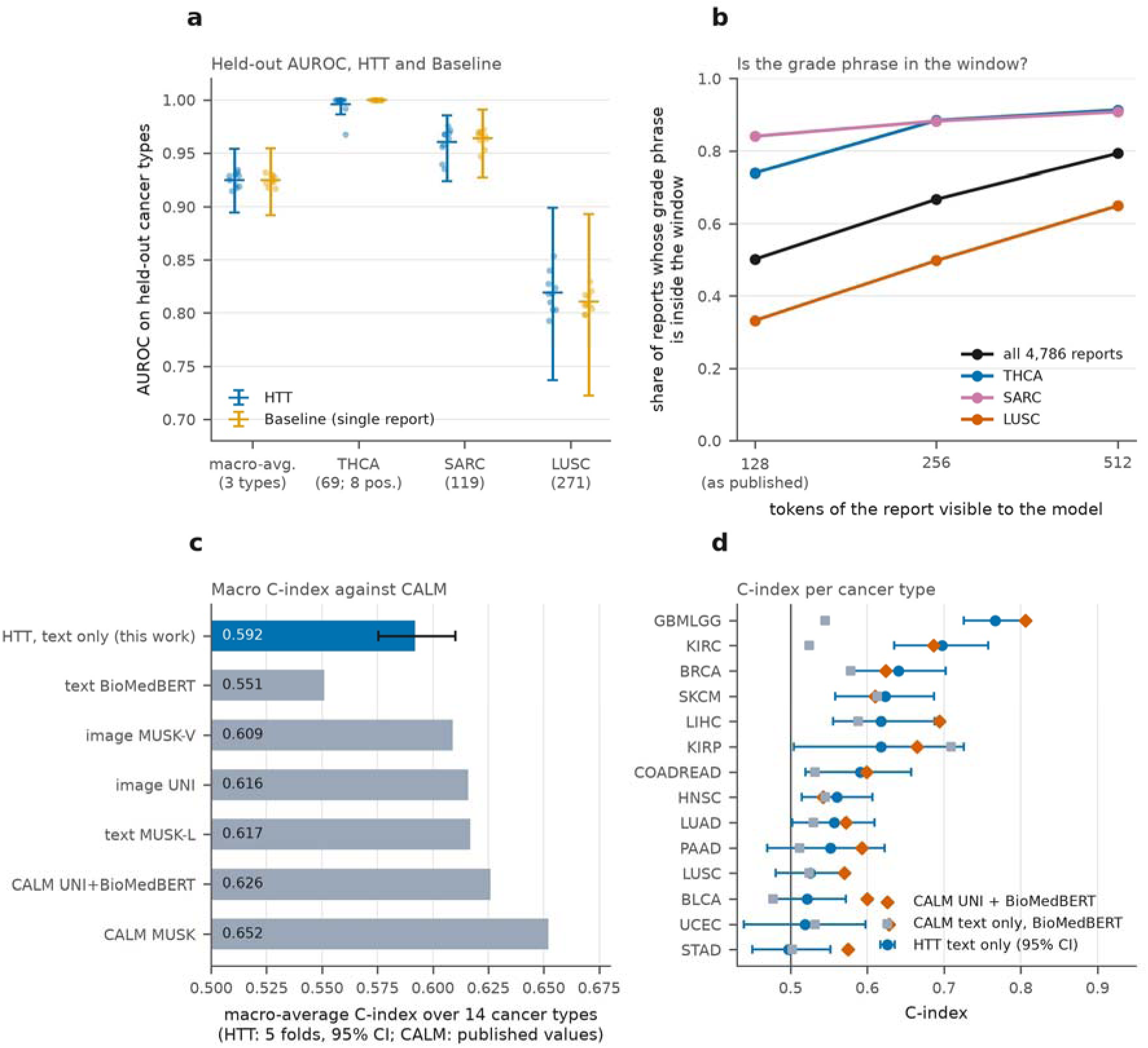
TCGA: held-out cancer types and comparison with CALM. (a) Held-out AUROC for HTT and the single-report Baseline: dots are seeds, whiskers the 95% class-stratified patient-bootstrap CI of the mean. (b) Share of reports in which the labelling rule finds the same grade phrase within the first 128, 256 and 512 tokens (128 is what the models read). (c) Macro-average C-index over 14 cancer types: text-only HTT (five folds, 95% CI) against CALM’s published values. (d) Per-type C-index: HTT with 95% CI against two CALM columns.

### 3.7 The pre-trained encoder carries the transfer

HTT does not beat its single-report Baseline on the withheld types: the macro difference is +0.000 (95% CI −0.008 to +0.009), and on the pooled held-out reports the Baseline is higher (−0.010, −0.018 to −0.002; 0 of 10 seeds). Because each patient has a single report, the temporal layer has nothing to order, and held-out types receive the reserved unseen-type index in place of a trained embedding, so type conditioning supplies little signal precisely where transfer is measured. Both facts bound what Level 2 can contribute in this experiment, and the comparison shows that the bound is binding. The transfer is therefore carried by the pre-trained encoder and the grade objective. BiomedBERT was pretrained on a corpus in which descriptors such as ‘poorly differentiated’, ‘high mitotic rate’ and ‘marked nuclear pleomorphism’ occupy neighbouring regions of representation space regardless of organ; fine-tuning maps that shared region onto the grade objective, and the mapping holds for unseen types whose descriptors fall within it.

Lung squamous carcinoma is the most informative case. It is the only withheld type whose anatomical site is represented in training, by lung adenocarcinoma, and shared vocabulary might have been expected to help; the opposite is observed (0.819, with lung adenocarcinoma also the weakest in-distribution type at 0.710). Two explanations are consistent with the data. One is representational interference: exposure to lung adenocarcinoma may induce a lung-specific encoding of grade language that is partly misaligned with squamous criteria, which rest on keratinisation and intercellular bridging rather than glandular architecture. The other is visibility of the label. The labelling rule finds the grade phrase within the first 128 tokens, the window read by the models, for only 50% of all reports and for 33% of lung squamous reports (74% for thyroid and 84% for sarcoma; Fig. 10b; Appendix Table A4), so part of the AUROC measures whether the phrase falls inside the window. We tested neither explanation, and distinguishing them would require a 512-token re-run and a design that withholds histological variants of trained sites alongside unrelated types.

### 3.8 A text-only HTT sits between the text branches of CALM

Trained on the PORPOISE cohort with CALM’s loss, bins and stopping rule, the text-only HTT reached a macro-average C-index of 0.592 (95% CI 0.575 to 0.610; Table 12; Fig. 10c,d). That is below full CALM with UNI (0.626) and with MUSK (0.652), above its BioMedBERT text-only branch (0.551; +0.041, 0.025 to 0.059; higher in 11 of 14 types) and below its MUSK-L text branch (0.617). The comparison is not like for like. The input differs (raw text against LLM-written summaries), the training recipe differs in learning rate and optimiser, CALM values are published and were not re-run, and part of our score comes from whether a report exists at all: in melanoma, where only 41% of patients have a report, the C-index is 0.623 on all patients and 0.461 on patients with a report. Per-type values are in Appendix Table A11.

**Table 12.**
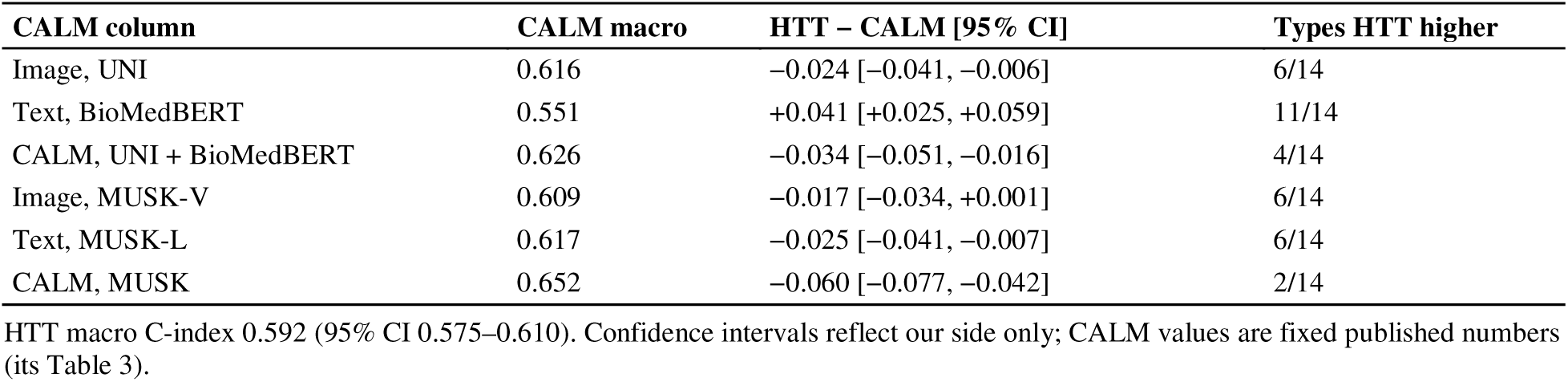
Macro-average C-index: text-only HTT minus each CALM column.

| **CALM column** | **CALM macro** | **HTT − CALM [95% CI]** | **Types HTT higher** |
|---|---|---|---|
| Image, UNI | 0.616 | −0.024 [−0.041, −0.006] | 6/14 |
| Text, BioMedBERT | 0.551 | +0.041 [+0.025, +0.059] | 11/14 |
| CALM, UNI + BioMedBERT | 0.626 | −0.034 [−0.051, −0.016] | 4/14 |
| Image, MUSK-V | 0.609 | −0.017 [−0.034, +0.001] | 6/14 |
| Text, MUSK-L | 0.617 | −0.025 [−0.041, −0.007] | 6/14 |
| CALM, MUSK | 0.652 | −0.060 [−0.077, −0.042] | 2/14 |

HTT macro C-index 0.592 (95% CI 0.575–0.610). Confidence intervals reflect our side only; CALM values are fixed published numbers (its Table 3).

## 4. Discussion

### 4.1 Principal findings

We developed a hierarchical, type-conditioned transformer for sequences of cancer reports and asked what each of its parts contributes, using benchmarks whose ceilings are known and enough seeds and patients to resolve the effects. Four findings stand out. Reading the whole sequence helps only modestly where a single report already carries most of the label, which is the case on the phrase benchmark, where the ceiling gap is 0.029. Visit timing cannot help in that regime, and the initial time code did not use it where it could have helped, on the growth benchmark. A clinical-scale time code that presents the interval to the previous visit makes the model respond to visit spacing and gains 0.053 AUROC, although it recovers only part of what the dates allow. On real reports, grade transfers to tumour types never seen in training at the level of in-distribution performance, and this is carried by the pre-trained encoder. Table 13 summarises the evidence by question.

**Table 13.** Summary of findings by question.

| Question | Finding | Key numbers |
|---|---|---|
| Q1. Does reading the sequence beat the last report? | Modestly on the phrase benchmark; most room is on unseen types | test180 +0.018 [−0.009, +0.050]; PANC100 +0.032 [+0.002, +0.070]; ceiling gap 0.029 |
| Q2a. Does timing help where it cannot carry information? | No; visit order speeds learning, real intervals add nothing | no time vs full −0.001 [−0.009, +0.007]; order only matches full at epoch 7 |
| Q2b. Does the initial time code use dates when they decide the label? | No | growth test180: 0.790 vs 0.787 without time; prediction shift 0.010 vs 0.374 |
| Q2c. Does a clinical-scale, interval-based code (HTT-IA) use them? | Yes, partly | 0.843 (+0.053 [+0.026, +0.082]); shift 0.273; below ceilings 0.894 and 0.979 |
| What makes learning fast under the protocol? | The per-visit auxiliary loss | 0/10 seeds learn without it on the phrase benchmark; take-off 7.9 vs 6.4 epochs on growth |
| Q3a. Does grade transfer to unseen tumour types? | Yes, on this task | 0.925 ± 0.007 [0.894, 0.954] vs in-distribution 0.928 ± 0.006 |
| Q3b. What carries the transfer? | The pre-trained encoder | HTT − Baseline +0.000 [−0.008, +0.009]; one report per patient |
| Q4. How does text-only HTT compare with CALM? | Between its text branches; below multimodal CALM | macro C-index 0.592 vs 0.551, 0.617, 0.626 and 0.652; inputs differ |

### 4.2 What a benchmark has to contain for timing to matter

The phrase benchmark is a sound test of whether a hierarchical model can aggregate evidence across reports, and in that respect it is informative: all reports raise the ceiling by at most 0.029 and HTT sits 0.030 below it. It cannot, however, show that a model uses timing, because the label is written into the text and the dates are generated by a process that does not touch it. For such data, the ceiling does not move when dates are added, and a model that learns to ignore them is behaving optimally. The growth benchmark shows what a benchmark that can answer the question has to look like. Dates are uninformative alone and informative through their interaction with a measured quantity (equation 2), so the model has to combine a number read from text with a timestamp. This corresponds to a common oncology situation in which lesion measurements are reported repeatedly and the meaning of a change depends on the interval. We do not claim that equation (2) is how real reports encode timing: free-text reports often contain dates, relative expressions such as ‘three months ago’, and qualitative statements that also constrain rate. The point is that identifiability can be built and verified, and that the ceiling gap with and without dates (Fig. 5) tells a reader whether a benchmark deserves to be used for this purpose.

### 4.3 Why the interval helps

The re-dating probe and the learned frequency scales suggest a coherent account, which we offer as an interpretation and not as a demonstrated cause. With frequencies between 1 and 0.05 rad per year and visits spread over under two years, the initial code behaves as a monotone ramp in elapsed time. The temporal transformer can read position from it but cannot distinguish a first follow-up at three weeks from one at three months, and the gradient reaching the frequency scales may be too small to change this, since they stayed at 1.004. The patient embedding does contain pace information (93% decodable on the phrase benchmark), so the encoding is not absent; it is not used because, there, the label does not require it. The advantage of the interval input over elapsed time is consistent with the structure of equation (2). Elapsed time requires the network to subtract two position codes to obtain the gap, whereas the interval presents the gap directly at the visit whose report contains the size change. The experiment that would separate this from an effect of the frequency range, the original frequencies with the interval input, was not run. The interval-only variant has no visit position and performed best, so ordering was not necessary on the growth benchmark, though it might be where order itself carries signal.

### 4.4 Why grade transfers across tumour types

The held-out result is non-trivial. A perfect AUROC on thyroid carcinoma, a type with its own grading vocabulary, anatomical context and histological features, shows that the model has not simply memorised type-specific keywords; all three withheld types were chosen because their grading language differs from everything in training, and all three transferred. The attribution experiment indicates where the capability lives. A generalisable representation of pathological aggressiveness appears to be formed during biomedical pretraining and recruited by fine-tuning [30], and progress on cross-type generalisation from single-report corpora depends primarily on encoder capability and label fidelity. The ordering of the held-out results runs against intuition, and as Section 3.7 explains it admits two explanations. If representational interference is real, selecting training types for a transfer-oriented system cannot proceed by maximising anatomical coverage alone, because a partly overlapping type may hurt performance on its histological neighbours.

### 4.5 Clinical framing

An AUROC of about 0.92 on unseen types supports assistive rather than autonomous use [31]. The in-distribution specificity of 0.62 reflects the class prior of the training corpus, and threshold calibration against the relative costs of over-and under-calling would precede any deployment. For triage the asymmetry is arguably the right one, since missing an aggressive tumour costs more than over-flagging an indolent one. Prioritisation is the defensible application: reports likely to describe aggressive disease are surfaced earlier for pathologist review, and the grade determination stays with the pathologist. Extension to staging, margin status or treatment response is not supported by the present experiments. Prevailing practice develops a separate model per cancer type; the transfer observed here indicates that a single model fine-tuned on a moderate number of common types can be applied to types absent from training, provided the target concept is already represented in the pre-trained encoder.

### 4.6 Recommendations for evaluating temporal clinical language models

The checks used here are inexpensive and none required access to proprietary data. We suggest that claims of temporal reasoning in longitudinal clinical text models be accompanied by:

a. a statement of how the label depends on timing in the data-generating process, or, for real data, evidence that timing is informative beyond order (for example, a model that sees only the dates);
b. ceilings for readers with and without the timing information, so that the available headroom is known before models are compared;
c. no-time and order-only ablations, and an intervention that re-dates fixed text, with the prediction shift compared with that of an informed reader;
d. baselines that receive the same supervision as the proposed model (here, the visit loss), and training long enough that a plateau is not mistaken for failure;
e. at least ten training seeds, a paired patient bootstrap rather than seed-level tests alone, and test sets large enough for the interval to exclude the effect sizes of interest, with checkpoints not selected on a set used as evidence.

These ideas are not new individually. They echo concerns about variance in machine-learning benchmarks [15], about shortcuts that let models appear to learn something they do not [32], and about leakage and over-optimism in applied machine-learning science [33]. Here they were jointly needed to establish what the temporal pathway of this architecture does, and the growth benchmark offers a small test bed with known ceilings for others.

### 4.7 Limitations

Everything except the TCGA experiments is synthetic, and the TCGA corpus has one report per patient, so the two capabilities are demonstrated on separate corpora. No result here shows HTT using timing on real longitudinal reports; that needs datasets with dated serial notes, such as MIMIC-IV or GENIE BPC, which we did not use. The synthetic results are a proof of mechanism under known ground truth and not an estimate of real-world performance. The growth benchmark is one construction: its growth model, noise level (8%), cadence range (14 to 180 days) and follow-up limit (365 days) were fixed before training and not varied, and the ceilings depend on them. Reports state sizes as numbers, which language models read imperfectly, and we did not separate a failure to read numbers from a failure to use time. The time-code variants are three variants with one setting each, evaluated at ten epochs and compared on the same test sets without correction for multiplicity; the frequency range and the input were changed together; and the interval-only variant discards order information that real data may need. HTT-IA was not evaluated on the phrase benchmark or at 20 epochs.

TCGA grade labels come from rule-based matching on the report text, which is standard and was used in the original TCGA-Reports benchmark [18] but admits label noise and, more consequentially, derives from the same text the model reads, so part of the performance may reflect recovery of matched phrases rather than generalisation. Two controls would bound this, a redacted condition replacing every matched span with a placeholder and a bag-of-words logistic regression on the same text, and expert review of a stratified subsample would quantify residual noise; none was done here, and only half of the labels are visible in the 128-token window. The three withheld types were chosen for diversity, not sampled systematically, so their mean should not be read as the expected performance on an arbitrary unseen type, and the sample sizes (69, 119 and 271 reports) leave wide intervals. Seeds vary training only, since the data split is fixed. Where a Wilcoxon test across seeds is significant and the bootstrap is not, we have said so and treated the bootstrap as the more cautious guide. The comparison with CALM uses published values for a different input, its metadata and refined splits are not public, and we ran one seed per fold.

All experiments use BiomedBERT-base under an 8 GB memory budget; larger encoders may yield stronger representations, particularly for rare types, and were left to future work.

## 5. Conclusion

The Hierarchical Temporal Transformer reads a patient’s sequence of cancer reports with a parameter-efficient report encoder, a cancer-type-conditioned temporal transformer and a dense per-visit loss. Evaluated against ceilings on controlled benchmarks, it benefits from reading the whole sequence where the sequence carries evidence, uses visit order from the first epochs, and uses real visit spacing only when the data make spacing informative and the time code is on a clinical scale, as in HTT-IA. On 4,786 real pathology reports it predicts tumour grade for tumour types never seen in training at the level of in-distribution performance, a capability that resides in the pre-trained encoder. Ceiling analysis, order-only and no-time ablations, and re-dating probes cost little and should accompany claims about temporal modelling. The next step is to test interval-aware modelling on real dated serial reports, where timing may enter the language of the notes as well as the metadata.

## Ethics statement

The study uses synthetic data and the public TCGA-Reports and PORPOISE resources; no new patient data were collected and no institutional approval was required.

## Acknowledgments

The authors thank the TCGA-Reports curators for releasing the corpus in machine-readable form.

## Data availability

TCGA-Reports [18] is openly available on the Hugging Face Hub. The model code, synthetic-data generator, label-extraction patterns, training configuration and evaluation scripts are available at https://github.com/rajneeshanand (OncoLocus, commit 47719d1). The growth-benchmark generator, ceiling and aggregation scripts, run-level outputs and figure scripts are [to be deposited in a public repository with a persistent identifier before submission].

## Appendix A. Supplementary tables

**Table A1a.** Phrase benchmark: trajectories.

| Trajectory | Prior | Visit label = 1 when | Patient positive? |
|---|---|---|---|
| Stable throughout | 0.30 | never | No |
| Progression midway | 0.30 | visit index ≥ □n / 2□ | Yes |
| Progression late | 0.25 | visit index ≥ n − 2 | Yes |
| Response then progress | 0.15 | last visit only, if visit index ≥ 2 | Yes, except when n = 2 |

Visit indices count from 0; n is the number of visits.

**Table A1b.** Phrase benchmark: evaluation sets.

| Set | Patients | Positive / negative | Composition |
|---|---|---|---|
| Validation | 60 | 36 / 24 | Published |
| test60 | 60 | 37 / 23 | Published test set |
| test120 | 120 | 73 / 47 | test60 + 15 extra per type |
| test180 | 180 | 113 / 67 | test60 + 30 extra per type (main test set) |
| PANC100 | 100 | 70 / 30 | Published zero-shot set |
| PANC300 | 300 | 199 / 101 | PANC100 + 200 extra |

Training and validation data are identical to the published run; extra patients come from the same generator with a disjoint seed (1042).

**Table A2.** Hyperparameters and training configuration (all experiments).

| Hyperparameter | Value | Note |
|---|---|---|
| Encoder | BiomedBERT-base-uncased | Biomedical pretraining; fits 8 GB memory |
| Max report length | 128 tokens | Longest synthetic report 59 tokens |
| Max sequence length | 10 visits (synthetic), 1 (TCGA) | Matches corpus structure |
| Hidden dimension; layers; heads; feed-forward | 512; 4; 8; 2,048 | Pre-LayerNorm |
| LoRA rank / alpha / dropout | 16 / 32 / 0.1 | Query and value projections; 11.1% trainable |
| Optimiser; learning rate; weight decay | AdamW; 2×10□□; 0.01 | Trainable parameters only |
| Warm-up | 10% of steps | From 10% of target rate |
| Batch; accumulation; gradient clipping | 4; 4 steps; 1.0 | Half-precision arithmetic |
| Auxiliary (visit) loss weight | 0.3 | Excluded at padded positions |
| Dropout | 0.1 (temporal), 0.2 (classifier) |  |
| Epochs; early stopping | 10; patience 5 on validation AUROC | 20 epochs without stopping for ablations |
| Seeds | 42 and 1–9 | Data split fixed at seed 42 |

All hyperparameters are shared across experiments; time-code variants change only the time input and the frequency range.

**Table A3.** Representative cancer-site extraction patterns for selected TCGA types.

| Code | Cancer type | Representative patterns |
|---|---|---|
| LUSC | Lung squamous cell | squamous.*lung, lung.*squamous |
| LUAD | Lung adenocarcinoma | \blung\b, pulmonary |
| BRCA | Breast cancer | \bbreast\b |
| KIRC | Kidney renal clear cell | \bkidney\b, renal cell |
| SARC | Sarcoma | leiomyosarcoma, \bsarcoma\b |
| THCA | Thyroid carcinoma | \bthyroid\b |
| GBM | Glioblastoma | glioblastoma, \bglioma\b |

Patterns are applied case-insensitively and most specific first; 17 ordered pattern sets were used in total.

**Table A4.** TCGA-Reports tumour-grade sets and visibility of the label in the first 128 tokens.

| **Set** | **Reports** | **Positive (high grade)** | **Negative** | **Median length (tokens)** | **Same label from 128 tokens** | **No grade phrase in window** |
|---|---|---|---|---|---|---|
| In-distribution test | 660 | 495 | 165 | – | – | – |
| THCA (held out) | 69 | 8 | 61 | 697 | 74% | 26% |
| SARC (held out) | 119 | 105 | 14 | 619 | 84% | 15% |
| LUSC (held out) | 271 | 246 | 25 | 991 | 33% | 66% |
| All labelled reports | 4,786 | – | – | 833 | 50% | 48% |

For all labelled reports, 2% of labels derived from the first 128 tokens differ from the full-report label.

**Table A5.**
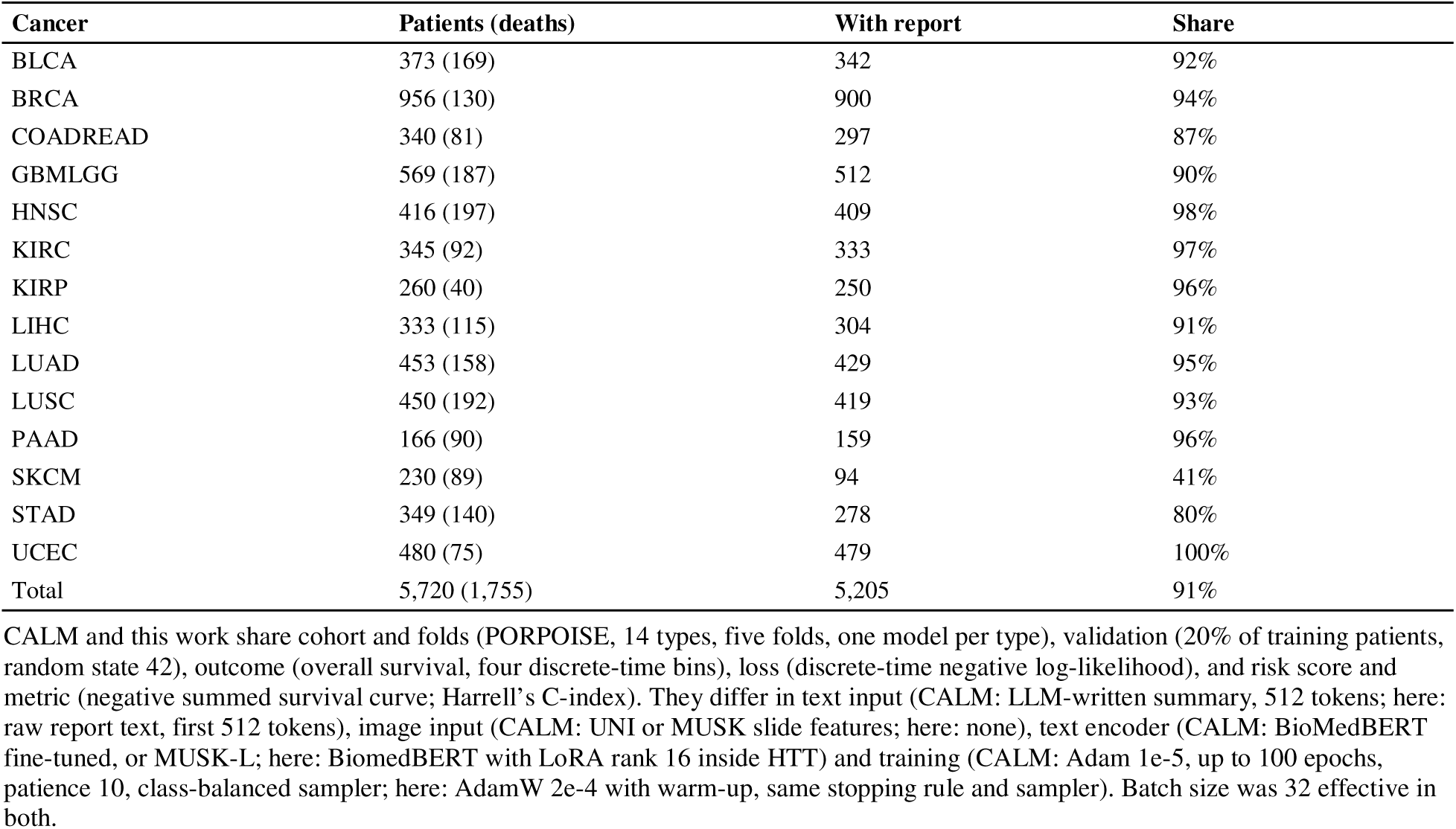
Survival cohort and protocol of the CALM comparison.

| Cancer | Patients (deaths) | With report | Share |
|---|---|---|---|
| BLCA | 373 (169) | 342 | 92% |
| BRCA | 956 (130) | 900 | 94% |
| COADREAD | 340 (81) | 297 | 87% |
| GBMLGG | 569 (187) | 512 | 90% |
| HNSC | 416 (197) | 409 | 98% |
| KIRC | 345 (92) | 333 | 97% |
| KIRP | 260 (40) | 250 | 96% |
| LIHC | 333 (115) | 304 | 91% |
| LUAD | 453 (158) | 429 | 95% |
| LUSC | 450 (192) | 419 | 93% |
| PAAD | 166 (90) | 159 | 96% |
| SKCM | 230 (89) | 94 | 41% |
| STAD | 349 (140) | 278 | 80% |
| UCEC | 480 (75) | 479 | 100% |
| Total | 5,720 (1,755) | 5,205 | 91% |

CALM and this work share cohort and folds (PORPOISE, 14 types, five folds, one model per type), validation (20% of training patients, random state 42), outcome (overall survival, four discrete-time bins), loss (discrete-time negative log-likelihood), and risk score and metric (negative summed survival curve; Harrell’s C-index). They differ in text input (CALM: LLM-written summary, 512 tokens; here: raw report text, first 512 tokens), image input (CALM: UNI or MUSK slide features; here: none), text encoder (CALM: BioMedBERT fine-tuned, or MUSK-L; here: BiomedBERT with LoRA rank 16 inside HTT) and training (CALM: Adam 1e-5, up to 100 epochs, patience 10, class-balanced sampler; here: AdamW 2e-4 with warm-up, same stopping rule and sampler). Batch size was 32 effective in both.

**Table A6.** Properties of the released phrase benchmark.

| **Property** | **Specification** | **Observed in regenerated data** |
|---|---|---|
| Trajectory mix | 30 / 30 / 25 / 15% | 31.8 / 30.4 / 23.2 / 14.6% (χ² p = 0.74) |
| Visits per patient | Uniform, 2–8 | 63–78 patients per count; mean 5.06 (χ² p = 0.93) |
| Patient positive rate | Not specified | 65.6% (code implies 67.9%) |
| Start year | 2018–2022 | 95–110 patients per year |
| Report length | Under 256 tokens | Maximum 59, mean 39 tokens; no truncation at 128 or 256 |
| Gap between visits | Day of visit k = k × uniform(42, 90), redrawn each visit | −225 to +371 days; 50% within 42–90; 10% ≤ 0; 35% of patients with out-of-order dates |
| Non-progressing phrases | Stable or improving wording | 50% stable, 50% improving; prostate has a change slot in 1 of 4 templates |

Regeneration reproduced split sizes and class counts exactly; file hashes were identical on two machines.

**Table A7.** Phrase benchmark: seed-42 run against the original run.

| **Metric** | **HTT published** | **HTT ours** | **Baseline published** | **Baseline ours** |
|---|---|---|---|---|
| Best validation AUROC | 0.942 | 0.939 | 0.881 | 0.873 |
| test60 AUROC | 0.912 | 0.927 | 0.937 | 0.934 |
| test60 F1 | 0.812 | 0.845 | 0.845 | 0.877 |
| PANC100 AUROC | 0.995 | 0.998 | 0.949 | 0.962 |
| PANC100 F1 | 0.947 | 0.978 | 0.882 | 0.889 |

The data are identical; differences come from hardware and library versions.

**Table A8.** Phrase benchmark, test180 AUROC by cancer type against the ceilings.

| **Cancer type** | **Baseline** | **Ceiling, last report** | **HTT** | **Ceiling, all reports** |
|---|---|---|---|---|
| All four types | 0.913 | 0.932 | 0.931 (0.937 at 20 epochs) | 0.961 |
| Breast | 1.000 | 1.000 | 0.997 | 1.000 |
| NSCLC | 0.967 | 0.975 | 0.956 | 0.997 |
| Colorectal | 0.926 | 0.905 | 0.929 | 0.978 |
| Prostate | 0.560 | 0.688 | 0.644 | 0.695 |

Mean of ten seeds. The baseline lies above its colorectal ceiling by chance: on 21 colorectal patients whose last report is uninformative it reacts to filler words that line up with the label in this sample.

**Table A9.** Phrase benchmark: what the patient embedding and time code encode.

| Metric | Value |
|---|---|
| Patient-embedding cosine, 21 vs 90 d | 0.985 |
| Embedding shift ÷ typical distance between patients | 0.13 |
| 21 vs 90 d decodable from the embedding | 93% (chance 50%) |
| Per-patient shifts aligned with one shared pace axis (mean cosine) | 0.68 |
| Position of 42 d and 66 d on that axis (21 d = 0, 90 d = 1) | 0.33 and 0.68 (linear in days: 0.30 and 0.65) |
| Cosine of pace axis with progressor − non-progressor axis | 0.89 |
| Pace shift ÷ progressor/non-progressor gap | 0.09 |
| Time-code cosine, 21 vs 90 d spacing, visit 1 / 3 / 5 / 7 | 0.997 / 0.973 / 0.926 / 0.860 |
| □ΔTE□ ÷ □report embedding□, visit 1 / 3 / 5 / 7 | 0.08 / 0.24 / 0.40 / 0.55 |

Ten published-protocol full models; the patient embedding is the classification-token output of the temporal transformer.

**Table A10.** Growth benchmark: initial time code trained for 20 epochs without early stopping.

| **Model** | **Validation** | **test60** | **test120** | **test180** | **PANC100** | **PANC300** |
|---|---|---|---|---|---|---|
| HTT, original time code | 0.904 ± 0.011 | 0.794 ± 0.030 | 0.831 ± 0.023 | 0.803 ± 0.023 | 0.826 ± 0.017 | 0.854 ± 0.015 |
| Baseline (last report) | 0.874 ± 0.005 | 0.717 ± 0.025 | 0.789 ± 0.019 | 0.775 ± 0.019 | 0.773 ± 0.020 | 0.827 ± 0.013 |
| HTT, no time input | 0.896 ± 0.005 | 0.782 ± 0.014 | 0.825 ± 0.015 | 0.798 ± 0.011 | 0.818 ± 0.007 | 0.850 ± 0.008 |

Mean ± SD over ten seeds. test180: HTT − baseline +0.029 [−0.022, +0.078] (7/10 seeds); HTT with and without time −0.006 [−0.013, +0.001]. The clinical-scale time codes were not run for 20 epochs.

**Table A11.** Text-only HTT against CALM: C-index per cancer type.

| **Cancer** | **Patients (events)** | **With report** | **HTT, mean ± SD over folds** | **CALM text BioMedBERT** | **CALM UNI + BioMedBERT** | **CALM text MUSK-L** | **CALM MUSK** |
|---|---|---|---|---|---|---|---|
| BLCA | 373 (169) | 342 | 0.522 ± 0.025 | 0.477 | 0.600 | 0.594 | 0.617 |
| BRCA | 956 (130) | 900 | 0.640 ± 0.065 | 0.578 | 0.624 | 0.598 | 0.606 |
| COADREAD | 340 (81) | 297 | 0.591 ± 0.074 | 0.532 | 0.599 | 0.636 | 0.666 |
| GBMLGG | 569 (187) | 512 | 0.767 ± 0.023 | 0.545 | 0.806 | 0.747 | 0.808 |
| HNSC | 416 (197) | 409 | 0.560 ± 0.041 | 0.545 | 0.542 | 0.561 | 0.567 |
| KIRC | 345 (92) | 333 | 0.697 ± 0.060 | 0.524 | 0.686 | 0.624 | 0.678 |
| KIRP | 260 (40) | 250 | 0.618 ± 0.199 | 0.709 | 0.665 | 0.740 | 0.756 |
| LIHC | 333 (115) | 304 | 0.618 ± 0.050 | 0.588 | 0.694 | 0.610 | 0.700 |
| LUAD | 453 (158) | 429 | 0.557 ± 0.070 | 0.530 | 0.572 | 0.614 | 0.629 |
| LUSC | 450 (192) | 419 | 0.526 ± 0.032 | 0.524 | 0.570 | 0.563 | 0.556 |
| PAAD | 166 (90) | 159 | 0.552 ± 0.088 | 0.512 | 0.593 | 0.494 | 0.584 |
| SKCM | 230 (89) | 94 | 0.623 ± 0.076 | 0.613 | 0.610 | 0.603 | 0.640 |
| STAD | 349 (140) | 278 | 0.498 ± 0.049 | 0.502 | 0.575 | 0.562 | 0.610 |
| UCEC | 480 (75) | 479 | 0.519 ± 0.116 | 0.532 | 0.628 | 0.689 | 0.713 |
| **Macro-average** |  |  | **0.592 [0.575, 0.610]** | **0.551** | **0.626** | **0.617** | **0.652** |

HTT: mean ± SD over five folds. CALM values are from its published Table 3.

